# An integrative trait-based framework to infer resource budgets and life-histories of long-lived plants

**DOI:** 10.1101/2023.04.29.538794

**Authors:** Huw Cooksley, Matthias Schleuning, Alexander Neu, Karen J. Esler, Frank M. Schurr

**Author notes:** Contact & corresponding author: Huw Cooksley –.

## Abstract

A fundamental assumption of functional ecology is that functional traits determine life-histories. Yet correlations between traits and life-history components are often weak, especially for long lived plants. This is because trade-offs, constraints, dynamic resource budgets and the scaling from single organs to entire plants cause complex relationships between traits and life-history. To elucidate these relationships, we present an integrated Trait-Resource-Life-History (TRL) framework that infers how functional traits affect organ-level costs and benefits of different life history components, how these costs and benefits shape the dynamics of whole-plant resource acquisition and allocation, and how these dynamics translate into life history. We illustrate this framework by developing a TRL model for a functionally diverse group of woody plants (22 species of the genus *Protea* from the South African Greater Cape Floristic Region). Using hierarchical Bayesian latent state-space modelling, we statistically parameterise this model from data on year-to-year variation in growth, reproduction and maternal care (serotiny) for 600 individuals. The parameterised model reveals that higher resource acquisition translates into both larger absolute resource pools and greater proportional resource allocation to reproduction. Accordingly, specific leaf area, a key trait increasing resource acquisition, is associated with larger resource pools, an earlier age of maturity as well as increased vegetative and reproductive performance at young to intermediate ages. In contrast, seed nitrogen content has opposing effects on the benefits of different organs and thus only shows weak correlations with life-history components. Importantly, the TRL model identifies trait and resource-mediated trade-offs at the level of organs, whole-plant resource budgets and life-histories. It can thus quantify key components of life-history theory that are so far largely inaccessible for long-lived plants. This permits novel insights into ecological and evolutionary mechanisms shaping life-histories. Application of the proposed framework to a broad range of plant systems should be facilitated by the increasing availability of trait and demographic data, whole-plant phenotyping and high resolution remote sensing. The integration of the TRL framework with models of biotic interactions further holds promise for a resource-based understanding of community dynamics across trophic levels and a closer integration of functional ecology, evolutionary ecology, community ecology and ecosystem science.

## Introduction

Understanding the drivers of organismal performance is foundational to ecology and evolutionary biology (Stearns, 1992). How individuals take up resources and allocate them to different dimensions of performance (such as growth or reproduction) shapes their life-histories, fitness and demographic rates, which in turn influence the dynamics of populations and communities (Shipley *et al*., 2016; Ohse *et al*., 2023). The vital rates of an individual, and its fitness in a given environment, hence arise from its resource budget.

Trait-based approaches have proliferated as a means to predict plant demography and fitness (Lavorel & Garnier, 2002; McGill, *et al*. 2006; Violle *et al*., 2014; Evans *et al*., 2016; Shipley *et al*., 2016; Salguero-Gómez *et al*., 2018; Laughlin *et al*., 2020). These approaches assume that functional traits influence the growth, survival, or reproductive rate of an individual (Violle *et al*., 2007; Reich, 2014; Yang *et al*., 2018). Despite the promise of trait-based approaches, numerous studies find perplexingly weak correlations between traits and whole-plant performance (Adler *et al*., 2013; Pistón *et al*., 2019; Swenson *et al*., 2020). This is especially the case in long-lived plants, which show substantial year-to-year variation in growth and reproduction and may experience trade-offs between performance early versus late in life (Petit & Hampe, 2006; Yang *et al*., 2018). Three challenges to a trait-based understanding of life histories are particularly pronounced in long-lived plants, such as trees and shrubs. These challenges are to understand how trait effects on resource costs and benefits scale up from organs to entire plants, how whole-plant costs and benefits shape resource allocation to different life-history dimensions, and how resource allocation changes through the lifetime of an individual. We expand on these three challenges in the following paragraphs.

Many functional traits are proxies for per-organ or per-unit resource investment and for the expected return on that investment. For instance, specific leaf area (SLA) reflects the costs of constructing and maintaining a given unit of leaf area, the expected energetic returns of this investment, and the susceptibility of leaves to biotic and abiotic stress (such as herbivory and desiccation) which reduce return on resource investment (Reich *et al*., 1997; Wright *et al*., 2004). However, since SLA is a measure per unit leaf mass, it has to be combined with whole-plant leaf biomass to determine plant-level construction costs and benefits in terms of carbon assimilation. Hence, it is necessary to scale up per-unit or per-organ traits to the whole plant level (Yang *et al*., 2018).

Constraints and trade-offs shape resource allocation in different ways (Stearns, 1992; Niklas, 1992). First, resource allocation is constrained by positive allometries if allocation to one organ must be matched by allocation to another. For instance, trees and shrubs cannot produce flowers and leaves without allocation to wood that provides mechanical support, water and nutrients (Niklas, 1992; 1994). Secondly, resource-mediated trade-offs constrain phenotypes if resource allocation to one organ comes at the expense of allocation to other organs, causing, for instance, a trade-off between reproductive and vegetative performance (Levins, 1968). Thirdly, trade-offs between the quantity and quality of organs can limit resource allocation (Moles & Westoby, 2006; Shipley *et al*., 2006). Such quantity-quality trade-offs arise if trait values that increase the costs of an organ (e.g. large seed size) also increase the organ’s benefits (e.g. higher germination and recruitment success). Identifying how traits and resources mediate these constraints and trade-offs is thus necessary to understand why organisms occupy a limited subset of the possible life-history space.

Changes in resource acquisition and allocation over an organism’s lifetime may furthermore lead to variation in trait-performance relationships with size and age (Iida *et al*., 2014; Gibert *et al*., 2016; Gray *et al*., 2019). For instance, the benefit of high SLA for vegetative growth may decrease with size in long-lived plants, as the shorter lifespan of high SLA leaves implies more investment into new support structures as plants grow (Gibert *et al*., 2016). As another example, plants with high seed mass or nutrient content should initially grow faster than ones with lower seed mass or nutrient content (Westoby, 1998; Cooksley *et al*., 2023). However, once they become reproductively mature, they need to invest more resources to produce a given number of seeds hence they may grow more slowly as adults and reproduce later (Stearns, 1992). Such size and/or age-dependent changes may thus contribute to weakening correlations between traits and whole-plant performance.

Quantifying how life-histories arise from trait effects on whole-plant costs and benefits of organs, from constraints and trade-offs governing resource budgets, and from age-dependent shifts in resource allocation should better elucidate the true relationships between traits and organismal performance and thereby improve the mechanistic understanding of life-history variation (Kleyer & Minden, 2015; Swenson *et al*., 2020). While this research agenda is highly promising, it faces considerable methodological challenges. First, it requires the quantification of variables that are very difficult to observe (such as whole-plant costs and benefits and resource budgets). Secondly, relationships between traits, whole-plant resource budgets and whole-plant performance are likely to be non-linear. A promising approach to overcome these challenges are latent state-space models (Clark, 2005). Latent state-space models can infer unobservable states and incorporate non-linear functional relationships resulting from physiological and ecological principles (such as allometric relationships or the principle of mass conservation; Enquist *et al*., 1999).

Here, we develop a Trait-Resource-Life-History (TRL) framework for latent state-space models to infer how functional traits shape whole-plant resource budgets and life-histories. In the following, we first outline the general concept of this framework. We then apply the TRL framework to the genus *Protea* from the South African Greater Cape Floristic Region (GCFR), a global hotspot of woody plant diversity (Myers *et al*., 2000; Born *et al*., 2007). Members of this genus exhibit a remarkable amount of variation in functional traits and life-history strategies (Schurr *et al*., 2012) and are well suited to elucidate long-term dynamics of resource budgets and life-history components. To this end, we develop a hierarchical Bayesian latent state-space version of the TRL framework that we parameterise from data on the life-history schedules of 600 *Protea* individuals. We use this empirically-parameterised TRL model to address three key questions of functional ecology and life-history theory: (1) do functional trait values that increase the benefits of resource investment into a life-history component also increase the resource costs of this investment? (2) What are the relationships between organ-level costs and benefits, whole-plant resource budgets and whole-plant performance, and do these relationships change with plant age? (3) Can functional traits be used to predict whole-plant performance and key life history characteristics? Building upon the *Protea* example, we then discuss the value of the TRL framework for an in-depth understanding of plant life-histories, the application of the framework to other study systems and the potential for its integration into process-based models of community dynamics across trophic levels.

## A Trait-Resource-Life-history (TRL) framework

The general structure of the TRL framework (Fig. 1) reflects a fundamental assumption of life history theory: in a given time interval, an individual acquires resources and can then allocate them to different life-history components such as vegetative growth, reproduction, parental care, survival and future resource acquisition (Stearns, 1992). Traits play a central role in the TRL framework as they determine the time-invariant resource costs and performance benefits of investment into a life-history component (Fig. 1). Since traits measure these costs and benefits at organ level, an implementation of the TRL framework has to track the amount of different organs per plant in order to quantify plant-level resource budgets and convert these into whole-plant life-history components (Fig. 1).

**Figure 1:**
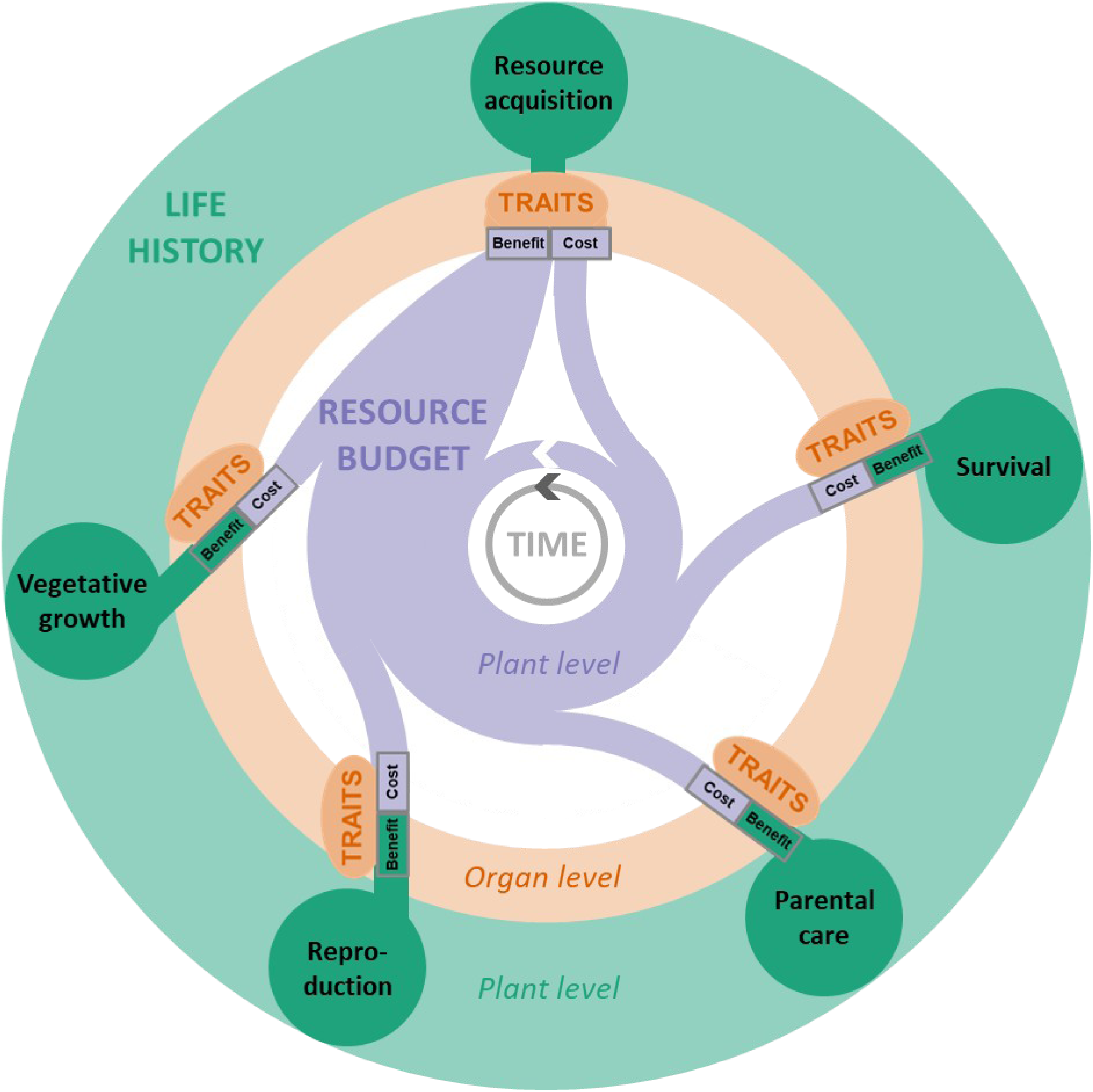
A trait-resource-life-history (TRL) framework for quantifying how functional traits shape resource budgets and life-histories of plants. Functional traits (orange ovals) shape organ level costs and benefits of resource investment into different life-history components (green circles). The framework requires quantifying how these functional traits scale up to whole-plant resource flows and how the resulting resource budget (purple) causes time-dependent changes of life-history components at the whole-plant level. While direct measurements of resource budgets are difficult, a latent-state-space representation of the TRL framework can be fitted to data on functional traits and life-history components (ovals and circles, respectively) and can model resource budgets as hidden aggregate variables.

Latent state-space modelling enables one to link this general framework to multiple types of data (Clark, 2005). In general, measurements of functional traits and proxies for life-history components are readily available whereas direct measurements of resource budgets are more difficult to obtain. In many cases, the resource will thus be modeled as a hidden aggregate variable that mediates trade-offs between life-history components rather than explicitly describing amounts of a single compound (Tonnabel *et al*., 2012). Modeling resource acquisition as a function of the amount of resource-capturing organs (e.g. leaves or roots) and expressing resource amounts in units of an easily measurable life-history component (e.g. wood biomass production), helps to constrain the model to biologically plausible parameter ranges and to improve its statistical identifiability.

## Application of the TRL framework to *Protea* plants

We applied the TRL framework to 20 species and 2 interspecific hybrids of the genus *Protea* (Proteaceae). For simplicity, all study taxa are referred to as ‘species’. They are endemic to the South African Greater Cape Floristic Region and frequently dominate fire-prone fynbos shrublands in this global biodiversity hotspot (Rebelo, 2001). These study species share the same fire-driven life cycle and general growth architecture. Their regular growth pattern and the fact that they form canopy seed banks enables one to reconstruct the temporal dynamics of growth, reproduction and maternal care for multiple years into the past (Walter *et al*., 2023). Despite this commonality, *Protea* species exhibit remarkable variation in life-histories and functional traits (Schurr *et al*., 2012), with adult plant height and seed mass varying by more than an order of magnitude (Treurnicht *et al*., 2016). In fact, Linnaeus named the genus *Protea* after the deity Proteus who – according to Greek mythology – could take any form. *Protea* species are thus well suited to test the TRL framework.

All study species are serotinous nonsprouters: fire kills them as adults but triggers the release of seeds from fire-protected infructescences (hereafter cones) borne in the adult canopy (Bond *et al*., 1984; Lamont *et al*., 1991). Almost all seeds then germinate in the first year after fire once soil moisture and temperature are conducive (Lamont & Groom, 1998). Consequently, our study species form even-aged stands. Within the first few years, *Protea* plants develop a deep taproot that accesses reliable water supply in deeper soil layers (Lamont & Groom, 1998; Cramer et al., 2014). Thereafter, they are very likely to survive until the next fire, unless this fire arrives exceptionally late (Bond, 1980; West *et al*., 2012; Treurnicht *et al*., 2016). Growth and survival during the critical juvenile phase increase with seed nutrient reserves (Cooksley *et al*., 2023).

*Protea* individuals show a regular growth pattern (Fig. 1) that is driven by strong seasonality, in particular in terms of rainfall seasonality in the Mediterranean-type climate found across most of the region (Bradshaw & Cowling, 2014). The branch increments produced in a given growth bout are separated by branching points or by ‘nodes’ (branch segments with a high density of leaves that are well visible by dense leaf marks even after leaves are shed). Branches typically produce one increment per year so that the number of increments along major growth axes can be used to age plants (Bond, 1985; Carlson *et al*., 2011; Treurnicht *et al*., 2016) although individuals growing in very benign conditions can consistently show either two or three branch increments per year. This regular growth pattern makes it possible to age individual branch increments and the subtending leaves, inflorescences and cones (Fig. 2; Walter *et al*., 2023). Moreover, it enables the reconstruction of past vegetative growth (see *Reconstructing whole plant vegetative growth*).

**Figure 2:**
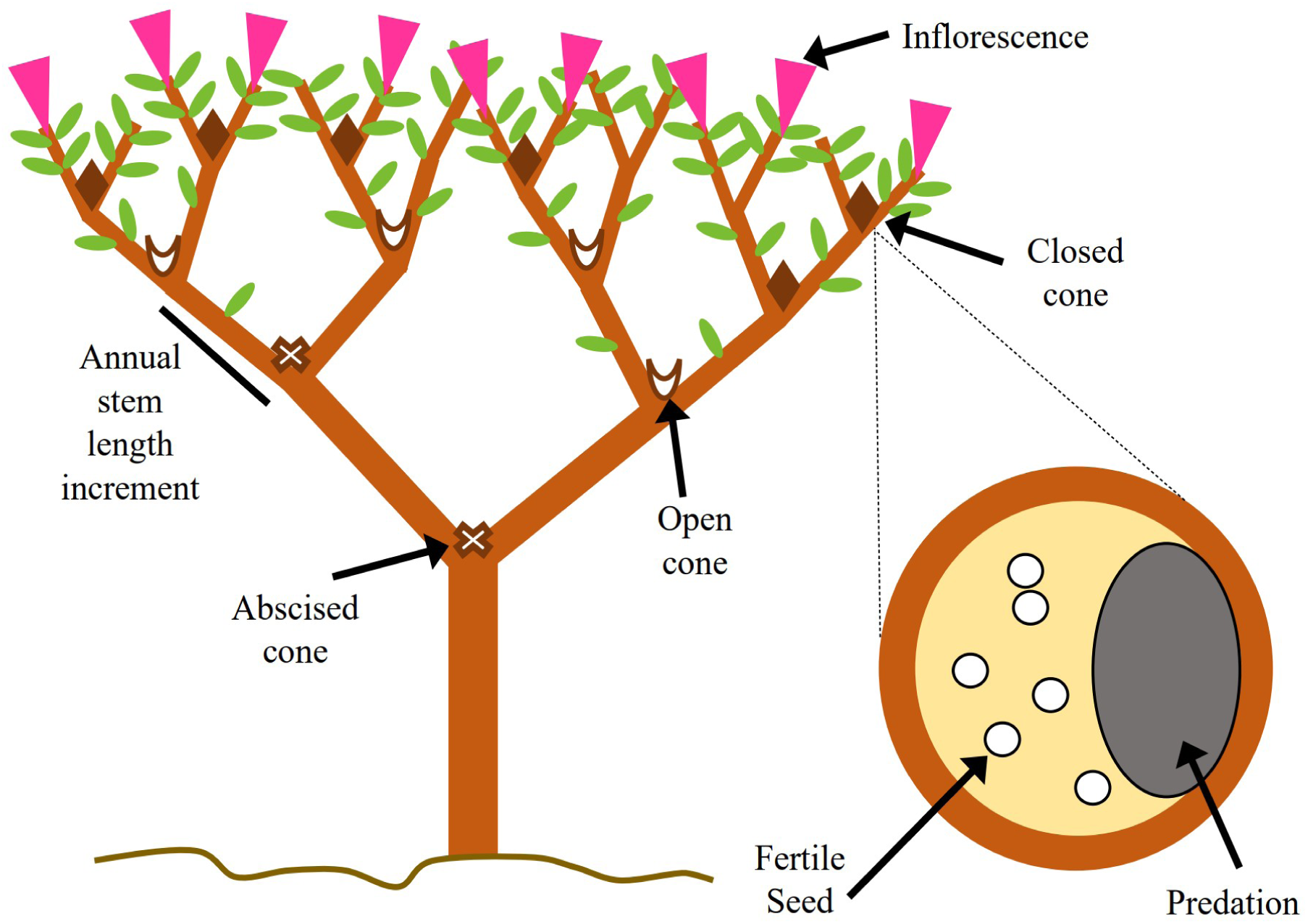
A stylised depiction of a *Protea* individual, showing proxies for life-history components measured in our study: stem length increments, and number and age of closed, open and abscised cones. The insert shows a stylised cross-section through a closed cone, showing fertile seeds and the proportion of the cone area consumed by insect seed predators.

*Protea* species are evergreen, retaining their sclerophyllous leaves for several years (Rebelo, 2001). While light typically does not limit plant growth in fynbos ecosystems (Cramer *et al*., 2014), Proteaceae are capable of using photosynthates to acquire limiting soil nutrients (notably P and N). They achieve this by producing root clusters (or proteoid roots) consisting of a high density of rootlets that actively excrete large amounts of carboxylates and phosphatases to mobilize inorganic and organic P (Shane & Lambers, 2005; Cramer *et al*., 2014; Lambers *et al*., 2015). Proteaceae can thereby effectively convert light into the acquisition of limiting soil nutrients.

Reproductive maturity is reached after several years but the study species differ considerably in age of maturity (Midgley & Rebelo, 2008). *Protea* plants are hermaphroditic, producing robust inflorescences that contain multiple florets (again with substantial interspecific variation in inflorescence size and floret number; Rebelo, 2001). Each floret contains a single ovule that can develop into a single-seeded fruit. Inflorescences produce copious amounts of nectar (Nottebrock *et al*., 2017) and are pollinated by a large diversity of insects and birds (Neu *et al*., 2023a).

After flowering, inflorescences turn into woody cones that can remain closed for several years (Bond, 1985). Our study species differ in degree of serotiny, which varies from weak to moderate (in comparison to strongly serotinous Australian Proteaceae, Lamont & Groom, 1998). Proteaceae cones need to be supplied with water and carbon to remain closed. In fact, closed cones continuously lose water and CO_2_ so that cone maintenance implies costs (Cramer & Midgley, 2009). Consequently, serotiny constitutes a case of maternal care in plants (Harris & Pannell, 2010; Tonnabel *et al*., 2012). Cones open when the water supply to them stops because the supporting branch died, because seed predators damaged xylem vessels or because the mother plant was killed in a fire. Open cones remain on the plant for many years after, and scars of abscised cones remain visible. Seedlings germinating from seeds released between fires face intense competition from established vegetation and have low recruitment success (Lamont *et al*., 1991). Moreover, our study species do not form persistent soil seed banks. Seeds that are released from the canopy seed bank after fire and germinate in competition-poor post-fire environments have relatively high establishment success. Consequently, the size of a plant’s canopy seed bank (all fertile seeds contained in closed cones) at the time of fire closely matches its lifetime reproductive output (Treurnicht *et al*., 2016).

The canopy seed banks formed by *Protea* species hold a substantial proportion of the above ground nutrient pool in fynbos ecosystems (in particular for P, N and K; Low & Lamont, 1986). This nutrient concentration results from the high nutrient acquisition ability of *Protea* plants (see above) and makes their canopy seed banks attractive to insect seed predators that complete their larval development inside *Protea* cones (Nottebrock *et al*., 2017). Seed predator communities comprise a limited number of species (Coleoptera, Lepidoptera and Diptera), most of which infest many different *Protea* species (Neu *et al*., 2023a). The majority of the seed predators are borers that bite through the woody involucral bracts to enter and/or leave cones (Coetzee & Giliomee, 1987).

### Reconstructing whole-plant vegetative growth

The reconstruction of past whole-plant wood and leaf biomass of *Protea* individuals rests on four findings (Fig. 3): (1) our study species conform to da Vinci’s rule on the conservation of stem cross-sectional area (Leonardo’s notes 394, 395, Richter, 1970), (2) there are strong species specific allometric relationships between trunk basal area and total stem length, (3) the mass of leaves produced on a branch segment is proportional to branch cross-sectional area (as predicted by the pipe model theory, Shinozaki *et al*., 1964), and (4) leaf retention can be described by species-specific Weibull functions. In the following, we describe the evidence for these findings and show how they can be used to reconstruct time series of whole-plant wood and leaf biomass of *Protea* individuals.

**Figure 3:**
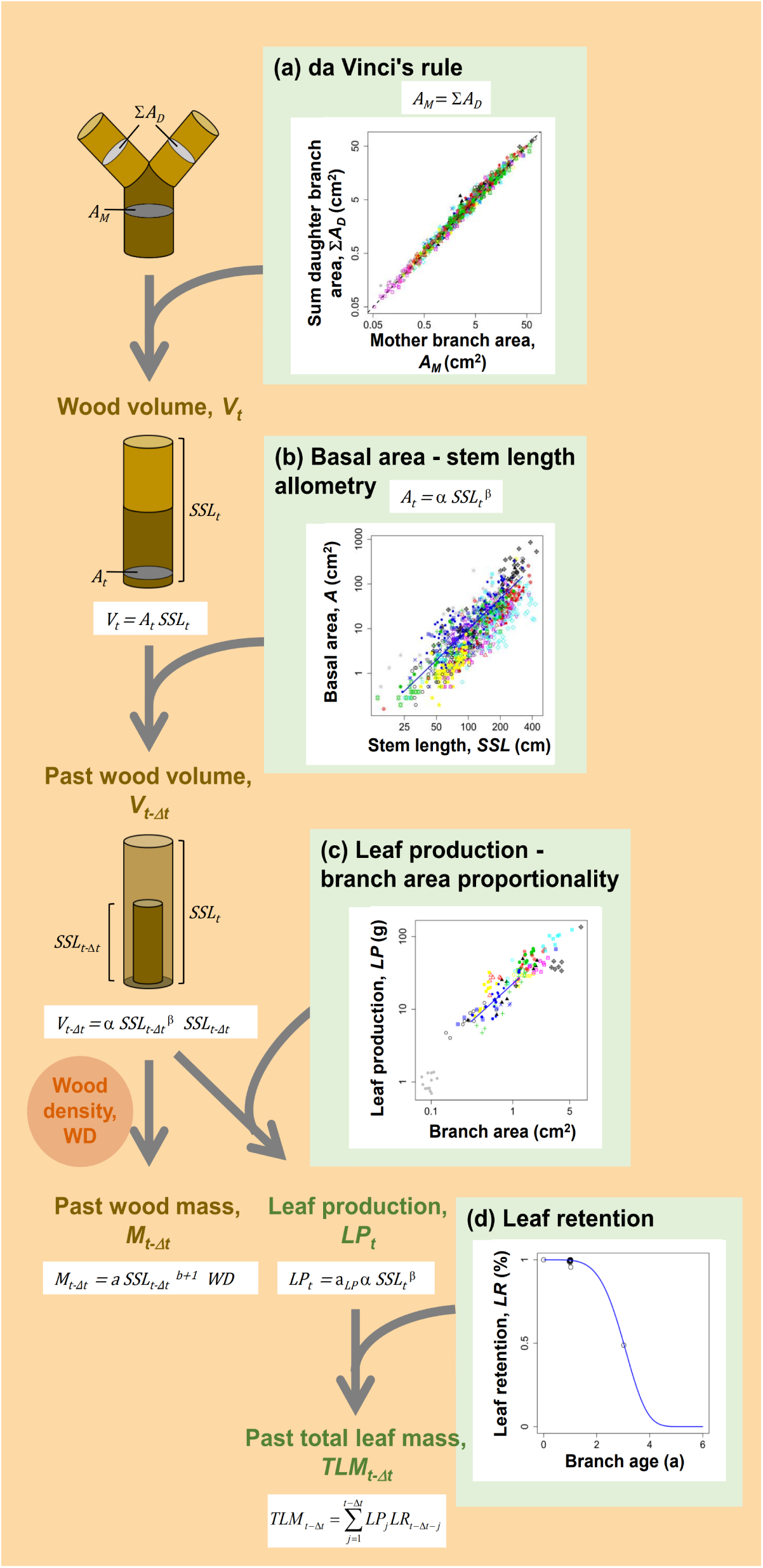
Reconstructing the vegetative growth of *Protea* individuals. (a) *Protea* plants follow da Vinci’s rule so that their wood volume can be approximated as the volume of a cylinder. (b) Since species-specific allometries (line of fit shown for *P*. *repens*) relate basal area to stem length, measurements of past stem length predict past wood volume and – in combination with wood density aboveground wood biomass in the past. (c) Species-specific proportionalities (line of fit shown for *P*. *repens*) between leaf mass produced on a branch and branch cross sectional area permit the prediction of whole-plant leaf production. By combining this with (d) species-specific leaf retention functions (line of fit shown for *P*. *repens*), one obtains proxies of whole-plant leaf mass in the past. For (a) – (c), species are shown as symbols of differing shape and colour.

Da Vinci’s rule states that the total cross-sectional area of branches is conserved across branching events (Richter, 1970), a prediction that is also derived from the pipe model theory (Shinozaki *et al*., 1964). To test whether this holds in *Protea*, we measured the diameter of the mother branch below a branching point and the diameters of all daughter branches above the branching point using callipers. We then calculated branch cross-sectional area as the area of a circle. This was done for 876 branching points across 425 individuals of all study species. We covered a wide range of branch sizes with mother branch cross-sectional areas ranging from 0.05 cm^2^ to 71.6 cm^2^. The obtained data show a very tight relationship between mother branch cross-sectional area (*A_M_*) and the summed cross-sectional areas of the daughter branches (Σ*A_D_*; Fig. 3a). In fact, a standardised major axis regression (SMA, Warton *et al*., 2007) relating log-transformed *A_M_* and Σ*A_D_* estimated a relationship that is very close to identity, with a power-law coefficient of 0.004 (95% confidence interval: -0.017 – 0.026) and a power-law exponent of 1.007 (95% confidence interval: 0.998 – 1.016). We thus find strong support for da Vinci’s rule in *Protea*. Consequently, the wood volume of a plant at time *t* can be approximated as the volume of a cylinder with basal area equal to trunk basal area *A_t_* and height equal to summed stem length *SSL_t_* (i.e. the sum of all stem length increments; Fig. 2).

To quantify allometric relationships between trunk basal area *A_t_*and summed stem length in the main growing direction *SSL_t_*, we measured these quantities for 1,041 plants of all study species (range of plant ages: 1-33 years). We then related log-transformed *A_t_* to log-transformed *SSL_t_* with a SMA regression including random effects of species identity on the intercept (Brose *et al*., 2019). This model of species-specific *A_t_ ∼ SSL_t_* allometries had a high goodness of fit (r^2^ = 0.85; Fig. 3B; Appendix S1: Figure S1). Consequently, trunk basal area at any time in the past *t - Δt* (A*_t-Δt_*) can be predicted from the observed summed stem length at this time point (*SSL_t-Δt_*). With the above-mentioned cylindrical approximation of wood volume, this enables the reconstruction of whole-plant wood volume at time *t - Δt* (Fig. 3). Multiplying this wood volume by population specific wood density, *WD*, yields whole-plant wood biomass as

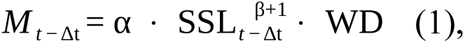

where α and β are the species-specific allometric coefficients and exponents, respectively, of the *A_t_ ∼ SSL_t_* – relationship. Consequently, wood biomass production in time interval *Δt* is

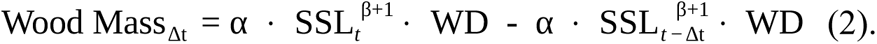

According to the pipe model theory (Shinozaki *et al*., 1964), the leaf mass produced on a branch is proportional to branch cross-sectional area. We tested this by estimating the allometric relationship between the leaf mass produced on a one-year-old branch and the branch’s cross sectional area. To this end, we collected all leaves from the branch and dried them at 70°C for 48 h before weighing them. In case leaves had been shed from the branch, we visually estimated the proportion of leaf mass lost (accounting for the loss of both entire leaves and leaf parts). This served to calculate the total dry mass of leaves produced on the branch. Additionally, we calculated branch cross-sectional area from calliper measurements of branch diameter.

Measurements on multiple branches per plant (mean: 2.7 branches) were then summed for each plant. We did this for 424 plants of all study species. A SMA regression of log-transformed total leaf mass and cross-sectional area with random effects of species and site on the intercept estimated a median allometric exponent of 1.0232 (CI: 0.9049 – 1.1459). Hence, there is a strong proportionality between leaf mass production and branch cross-sectional area (Fig. 3c; Appendix S1: Figure S2). In combination with da Vinci’s rule, the total leaf mass that a plant produces in year *t*, *LP_t_*, is thus proportional to the plant’s trunk basal area *A_t_* and to *SSL ^β^*

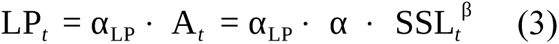

where *α _LP_* is the species-specific coefficient relating leaf mass production to branch cross sectional area.

Since *Protea* plants are evergreen, their standing leaf biomass depends not only on leaf production in a given year but also on leaf retention from previous years. To characterize leaf retention, we visually estimated the proportion of leaf mass retained on one year old branches of 458 individuals of all study species. Along the same branches, we estimated the branch age at which the proportion of retained leaf mass drops to 50% (cf. Treurnicht *et al*., 2020). These estimates of median leaf retention time were combined with corresponding measures from the FYNBASE database (Schurr *et al*., 2007; Treurnicht *et al*., 2020) in TRY (Kattge *et al*., 2020) to calculate species-level medians of median leaf retention time. We fitted species-specific Weibull functions to all leaf retention data for a species, since the Weibull distribution provides good descriptions of leaf lifespan in evergreen Mediterranean shrubs (Dungan *et al*., 2008).

Specifically, we used non-linear least squares to fit the complement of the Weibull’s cumulative density function, which describes the proportion of leaf mass retained on branches of age *k*, *LR_k_* as *LR =* exp¿, where *a* and *s* are the shape and scale parameter of the Weibull distribution, respectively. From the resulting species-specific leaf retention functions (Fig. 3d; Appendix S1: Figure S3), one can reconstruct a plant’s *total* standing leaf mass, *TLM*, at time *t - Δt* in the past as:

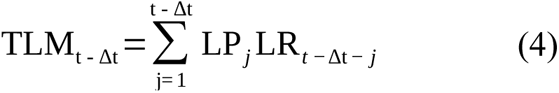

In summary, species-specific scaling relationships enable one to effectively reconstruct the vegetative growth of *Protea* plants. In particular, measurements of stem length increments in past growth bouts make it possible to calculate proxies of whole-plant aboveground wood biomass, leaf biomass production and total leaf biomass at multiple time points in the past (Fig. 3).

### Measurement of life-history components

We took time-stratified measures of vegetative growth, reproduction and maternal care for 600 individuals of the 22 *Protea* species on 21 study sites. By comparing the known time of last fire at each study site to the number of growth increments per focal plant (Bond, 1985; Treurnicht *et al*., 2016), we confirmed that the study individuals had established after the last fire. For 160 plants from five sites, the number of growth increments was either two or three-times larger than the site’s post-fire age, indicating that the plants had undergone two or three growth bouts per year, respectively. By combining known post-fire ages of each site and counts of growth increments, we were thus able to age both individual increments and entire plants.

To reconstruct the vegetative growth of each individual, we measured the length of each growth increment (the distance between nodes) along a representative stem in the plant’s main growing direction (starting from the youngest increment). For most individuals, these increments represent annual growth. However, in seeing two or three growth bouts per year, we took these measures at higher temporal resolution. In older stem sections, for which growth increments could not be delineated unambiguously, we additionally recorded the total stem length from the last resolvable increment to trunk base as a measure for total plant growth up to this time point.

Reproductive performance was quantified by cone production, retention of closed cones, seed set in sampled closed cones, and cone defence from predation. Cone production and closed cone retention was measured by counting the number of cones produced by each individual at each time-step (at the same temporal resolution as for growth increments), and whether they were closed, open or abscised at the time of observation. In certain older plants, it was not possible to assess reproduction in the first years of life; for these initial years, cone number was recorded as missing data. In very large plants, cones were counted in a representative subset of the canopy and these counts were then extrapolated to the entire plant. From each cohort of closed cones on an individual, we sampled one closed cone to measure seed set (seed number per cone). To this end, the cone was cross-sectioned at the stratum where seeds are borne, and the number of fertile seeds was counted (Treurnicht *et al*., 2016). In each cone, we also visually estimated cone defence as the proportion (in steps of 5%) of the total seed area not consumed by seed predators. In summary, our time-stratified measures of life-history components reached back 1-30 years, thus encompassing much of the total lifespan of our focal plants, which ranged from 3-33 years at the time of sampling.

### Measurement of functional traits

To quantify the costs and benefits of resource allocation to different life-history components, we measured four key functional traits representing investment into growth and reproduction, and return on this investment: specific leaf area (SLA), leaf N concentration, cone mass and seed N content (Appendix S1: Figure S4; Table S1). Additionally, we measured wood density of one year old stem wood as a key determinant of whole-plant wood biomass (see eqn. 1). For SLA, leaf N concentration and wood density, we followed the standard protocols of Pérez Harguindeguy *et al*. (2013; leaf N concentration was measured using elemental analysis (Vario EL Cube, Elementar Analysensysteme GmbH, Langenselbold, Germany)). Cone dry mass was measured from closed cones that were oven-dried at 70°C for 48 h and then weighed using precision scales. Seed dry mass (g) was measured from fertile seeds oven-dried at 70°C for 48 h, and weighed with precision scales. Seed N concentration (% mass) was determined by elemental analysis of finely milled dried seeds (Vario EL Cube, Elementar Analysensysteme GmbH, Langenselbold, Germany). Seed N content was then calculated as the product of seed mass and seed N concentration. In addition, we used measurements of leaf and seed N concentration from the FYNBASE database. Note that seed N and P contents are strongly correlated in these species (r = 0.86; Cooksley *et al*., 2023). Trait measurements were averaged at population level, except for leaf N which were averaged at species level.

### A Trait-Resource-Life-history (TRL) model for *Protea*

Based on the knowledge of the life cycle of our study species, we adapted the general TRL framework (Fig. 1) in four ways (Fig. 4). First, we did not explicitly model resource investment into survival (since all study species have very high rates of inter-fire survival). Secondly, we resolved reproduction into cone and seed production, maternal care in the form of cone maintenance and defence against seed predation. Thirdly, we modeled resource acquisition as a function of leaf mass because *Protea* plants are capable of converting photosynthates into acquisition of soil nutrients, water and pollination service (see above) and because leaf biomass can be reconstructed for past time points (see above) and is allometrically related to root biomass (Niklas, 2004). Fourthly, we expressed resource amounts in units of wood biomass which can also be reconstructed for our study species (see above), with conversion parameters for the relative costs of other tissues. In brief, this TRL model is an integration of organ-level costs and benefits informed by traits, whole-plant resource pools at each time-step, and proportional allocation of resources to wood, leaves, cones, seeds and serotiny, which we detail in the following.

**Figure 4:**
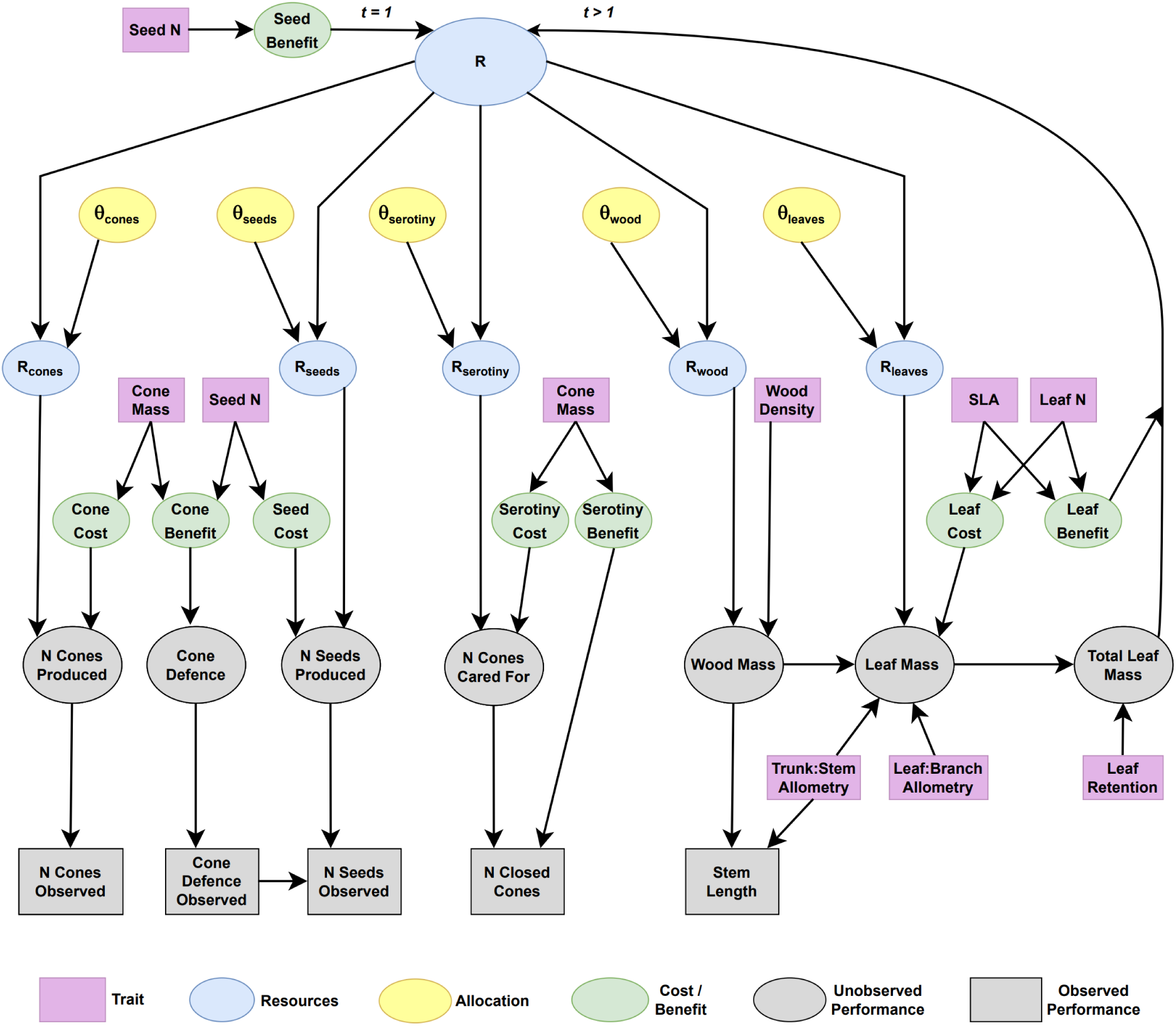
Process diagram of the TRL model for *Protea*. Data are represented by rectangles, and unobserved nodes are represented by ovals. Costs and benefits of different organs are informed by traits. In turn, organ costs, allocations and the resource pool determine the production of cones, seeds, wood and leaves, and maternal care. These latent performance measures are then linked to the observed performance data. Seed benefit determines the amount of resources available to a seedling in the first time-step, and for subsequent time-steps leaf benefit and total leaf mass determine resource acquisition and the size of the resource pool in the next time-step. Serotiny benefit determines the proportion of cones surviving to the next time-step.

Organ-level costs and benefits are defined with respect to resources (in units of equivalent grams of woody biomass). Costs of leaves, cones, seeds and serotiny are the resources required to produce one gram of leaves, one cone or seed, or to maintain one closed cone for one year, respectively. The benefits of leaves, cone defence, seeds and serotiny represent the respective return on these resource investments. Leaf benefit is the amount of resources acquired per gram of leaf and year. Seed benefit is the resource pool a seedling derives from the seed. Cone defence benefit is the per-year and per-cone-area probability of successful defence against seed predators. Serotiny benefit is the per-year probability of a cone remaining closed (and thus potentially contributing to post-fire regeneration).

Costs and benefits are modelled as functions of key traits for the respective organ, with random intercepts for species, site and individual (Table 1). While costs and benefits thus vary between individuals, we assume that they do not change with the age of an individual. Leaf cost and benefit are informed by SLA and leaf N concentration (Wright *et al.,* 2004). Cone cost is a function of cone dry mass, as the dense wood and bracts that make up most of a cone and afford protection for seeds are likely the substantial cost of this organ. Cone defence benefit is informed by cone dry mass, representing the cost to the parent plant of defending seeds, and by seed N content, representing the potential reward for granivorous insects. Seed cost and benefit are functions of seed N content, a key limiting nutrient in impoverished fynbos soils and a good predictor of juvenile performance (Cooksley *et al*., 2023). We model serotiny cost and benefit as functions of cone dry mass, assuming that costs of water replacement and carbon supply are proportional to cone dry mass (water content and dry mass of cones are highly correlated: Pearson’s r=0.87). Since resources are measured in units of wood biomass, we do not explicitly estimate the cost of wood production, but rather take it to be wood density.

**Table 1:** Overview of formulae for costs and benefits of resource allocation to life-history components.

| **Cost / Benefit** | **Link** | **Trait covariates** |
| --- | --- | --- |
| Leaf cost | ln – ln | SLA, Leaf N concentration |
| Cone cost | ln – ln | Cone mass |
| Seed cost | ln – ln | Seed N content |
| Serotiny cost | ln – ln | Cone mass |
| Leaf benefit | ln – ln | SLA, Leaf N concentration |
| Cone defence benefit | logit – ln | Cone mass, Seed N content |
| Seed benefit | ln – ln | Seed N content |
| Serotiny benefit | logit – ln | Cone mass |

All formulae include normally-distributed, zero-mean random intercepts for species, site and individual.

Whole-plant performance components of individual *i* in time-step *t* are latent variables modelled as a function of the resource pool, *R*, proportional resource allocations, *Θ*, to each component, and the respective organ-level cost. The performance variables are grams of wood and leaves produced, number of cones and seeds produced, as well as number of closed cones cared for.

We first model the reproductive maturity status, *Mature*, of a plant as a function of *R*

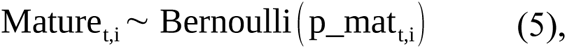

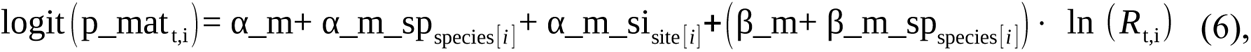

where *α_m_sp* and *α_m_si* are zero-mean normal random intercepts for species and site, respectively, and *β_m_sp* are zero-mean normal random slopes for species.

Proportional resource allocation to wood, leaf, cone and seed production and cone maintenance forms a five-element allocation vector ***Θ*** = (*Θ_W_*, *Θ_L_*, *Θ_C_*, *Θ_S_*, *Θ_M_*) defined by

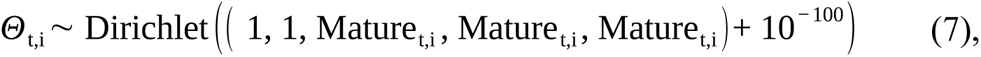

such that allocations are assigned a flat Dirichlet prior, except that allocations to cones, seeds and serotiny are essentially zero if an individual is immature. We thus assume that all resources in a given time-step are used, with no carry-over to the next time-step.

The five whole-plant latent performance variables are then defined by:

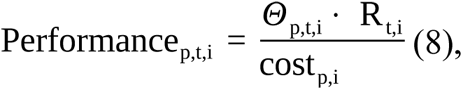

where *p* denotes the respective performance variable.

An individual’s resource pool in the first time step is a function of seed benefit

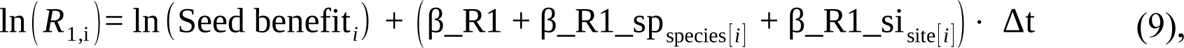

where *β_R1* and the zero-mean normal random slopes for species and site of *β_R1_sp* and *β_R1_si* define resource growth from seed resources to the resource pool in the first time-step, *Δ t* (1, 1/2 or 1/3 year, depending on the number of growth bouts per year, see above).

The resource pool of an individual in subsequent time-steps is a random variable, the expected value of which depends on leaf benefit and *TLM* in the current time-step. Thus:

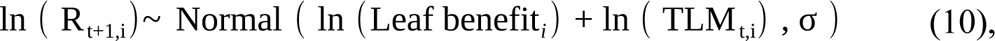

for all *t* >1, where *σ* is the standard deviation of the residual error.

The expected number of closed cones on a plant, *N closed cones*, is a function of *N cones cared for* and *Serotiny benefit*, assuming all closed cones on a plant in a given year receive the same allocation to serotiny

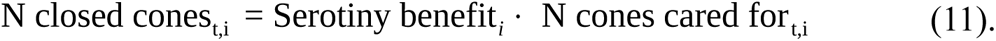

Cone defence against granivorous insects is defined as the probability, *p_def*, that a given 5% section of the total seed basal area is not eaten. This is modelled as a function of cone defence benefit (*Defence benefit*), site age at the time of cone production (*SA*, reflecting post-fire changes in predation; Walter *et al*., 2023) and cone age, *CA*, the time a cone has been on a plant before the year of observation:

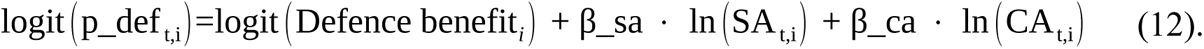

We link latent performance variables to age-stratified observations of performance via stochastic observation equations. The number of cones observed, *N cones obs*, was modelled as

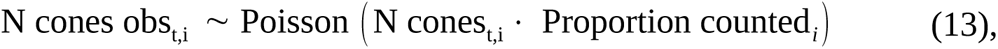

where *Proportion counted* is an offset for the proportion of an individual for which cones were counted. Observed cone defence, *Cone defence obs*, was modelled as

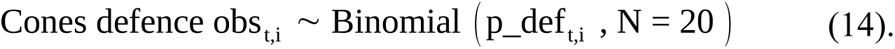

The number of seeds observed in a cone, *N seeds obs*, was modelled as a function of cone defence as well as whole-plant seed and cone production

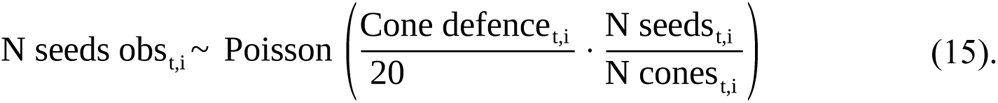

The total number of closed cones observed on a plant, *N closed cones obs*, was modelled as

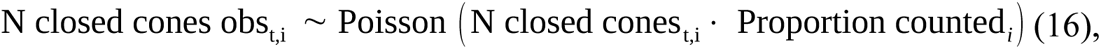

and the observed cone-age-stratified number of closed cones, *N closed cones obs str*, was

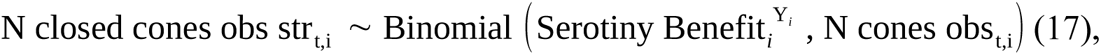

where *Y_i_* is the age of cones of that cohort in years. By rearranging equation 2, we obtain

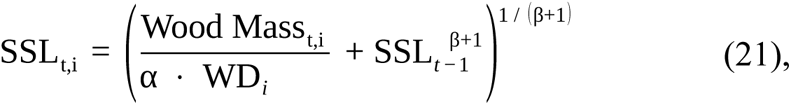

so that the expected stem length increment, *SL_t_*, is

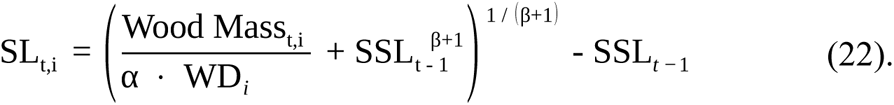

Since the allometry of seedlings differs from that of older plants (Niklas, 2004), we allowed the allometric relationship between *SSL_t_* and *A_t_* (Fig. 3) to differ between the first time-step and later time-steps. To this end, we estimated an α of seedlings that was multiplied by a species-specific random effect. Since in our study species, variance in stem length increments between branches of a given cohort is approximately equal to the mean (see Appendix S1: Figure S5), we modelled observed stem length increment as

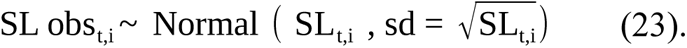

We informed resource-based estimates of leaf production with the proportional relationship between leaf mass and branch cross-sectional area by combining equations 3 and 9, such that

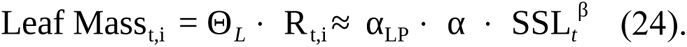

By ln-transforming, incorporating uncertainty and rearranging to maintain a proper acyclical graph, we obtain

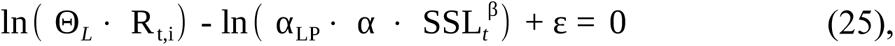

where *ε* is the residual error of this relationship, given an informative prior based on our empirical estimate (see ‘Reconstructing whole-plant vegetative growth’). Similar to above, we allowed the proportionality between leaf mass and cross sectional area to vary for plants in the first time-step, adding random intercepts for species and site to equation 25 for *t* = 1.

### Fitting the TRL model to data

All data preparation, calculations and statistical analyses were performed in R 4.0.0 (R Core Team, 2020). Our model was fitted to data via Markov chain Monte Carlo (MCMC) sampling, using JAGS 4.3.0 (Plummer, 2003) interfaced via the R package saveJAGS 0.0.4.9002 (Meredith, 2021). Three MCMC chains were run, with 20,000 iterations of adaptation and 40,000 iterations of sampling, thinned to every 20th iteration, resulting in 6,000 samples from the joint posterior distribution. We used weakly informative priors for all parameters, except for the effect of seed N content on seed cost, and for the residual error of leaf mass per branch area, which were informed by data for these processes. JAGS and R code used to fit the TRL model, including prior distributions, are given in Appendix S2. Convergence to the stationary posterior distribution was determined by the potential scale reduction factors for all process parameters being < 1.1, and by visual inspection of trace plots.

### Analysis of the parameterised model

Posterior predictions of observed performance measures were done as ‘one-step-ahead’ predictions, conditioned upon the previous resource state. In short, we simulated resource amounts from the whole-plant resource acquisition state-transition equation (equation 10) for the previous time step and then drew all other parameters from the joint posterior distribution.

To assess whether functional trait values that increase the benefits of resource investment into a life-history component also increase the resource costs of this investment, we compared posterior distributions of effect sizes of traits on paired costs and benefits.

To assess relationships between organ-level costs and benefits, whole-plant resource budgets and whole-plant performance, we calculated Pearson’s correlation coefficients between posterior medians of costs and benefits, resource pools, allocations and yearly whole-plant performance.

We conservatively chose to calculate these correlations beginning from the age of 6 years, as prior to this numerous posterior medians were zero, distorting correlations, and ending at 18 years, as few species had older individuals in the dataset. Prior to calculating correlation coefficients, costs and benefits were ln-transformed (except for cone and serotiny benefits which were logit-transformed), cone, seed and closed cone number were ln(x+1)-transformed, leaf and wood masses as well as traits were ln-transformed.

To address whether functional traits can predict whole-plant performance measures, we calculated Pearson’s correlation coefficients between them for ages from 6 to 18 years. We also calculated posterior median total reproductive output (seed bank size) for each individual at each time-step, integrating over cone and seed production, cone defence and cone maintenance, and calculated correlation coefficients of this fitness proxy with traits across the above age range.

Finally, we calculated Pearson’s correlations between traits and age of maturity (the earliest age at which the posterior median of seed bank size was ≥ 1).

## Results

### Predictions of life-history components and resource budgets

The TRL model predicted most performance measures with high accuracy and little bias (Fig. 5). For the four performance measures directly affected by resource allocation (stem length increment, cone number, seed number and closed cone number), the squared correlation coefficient (r^2^) between observed and posterior-median one-step-ahead predicted values ranged from 0.73 to 0.95 (Fig. 5a-d). Predictions of cone defence (which are not directly informed by resource budgets) were still reasonably accurate (r^2^ = 0.54) (Fig. 5e).

**Figure 5:**
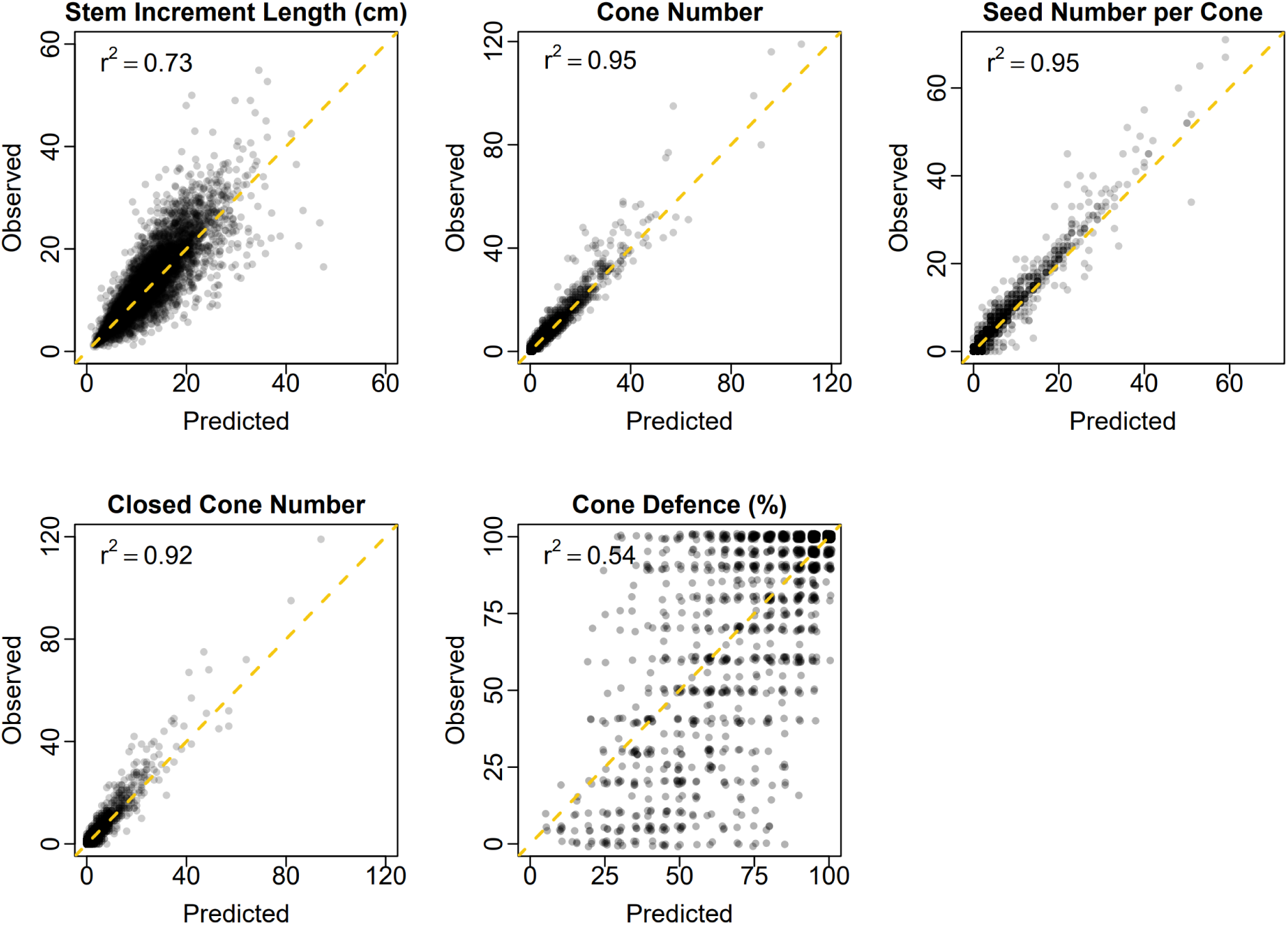
Observed annual performance measures for 600 *Protea* individuals against predictions of the TRL model for (a) annual stem length increment, (b) annual cone production, (c) seed number per sampled cone at time of observation, (d) number of closed cones maintained (age stratified) at time of observation and (e) cone defence percent at time of observation (points are jittered slightly for clarity). Predictions are posterior medians of one-step-ahead predictions conditioned upon the resource state in the previous time-step. The square of the Pearson’s correlation between observed and predicted values is shown in the top-left corner of each plot, and the dashed gold line represents the line of identity.

In addition to predicting observable performance components, the TRL model also estimates the essentially unobservable dynamics of whole-plant resource budgets (Fig. 6a). Specifically, it permits the individual and age-specific quantification of resource acquisition that leads to growth of the resource pool and of the allocation of this resource pool to different components (Fig. 6a). Moreover, the TRL estimates the build-up of total reproductive output (which for *Protea* individuals is measured as the size of the canopy seed bank; Fig. 6b) which integrates over cone and seed production, cone defence and serotiny.

**Figure 6:**
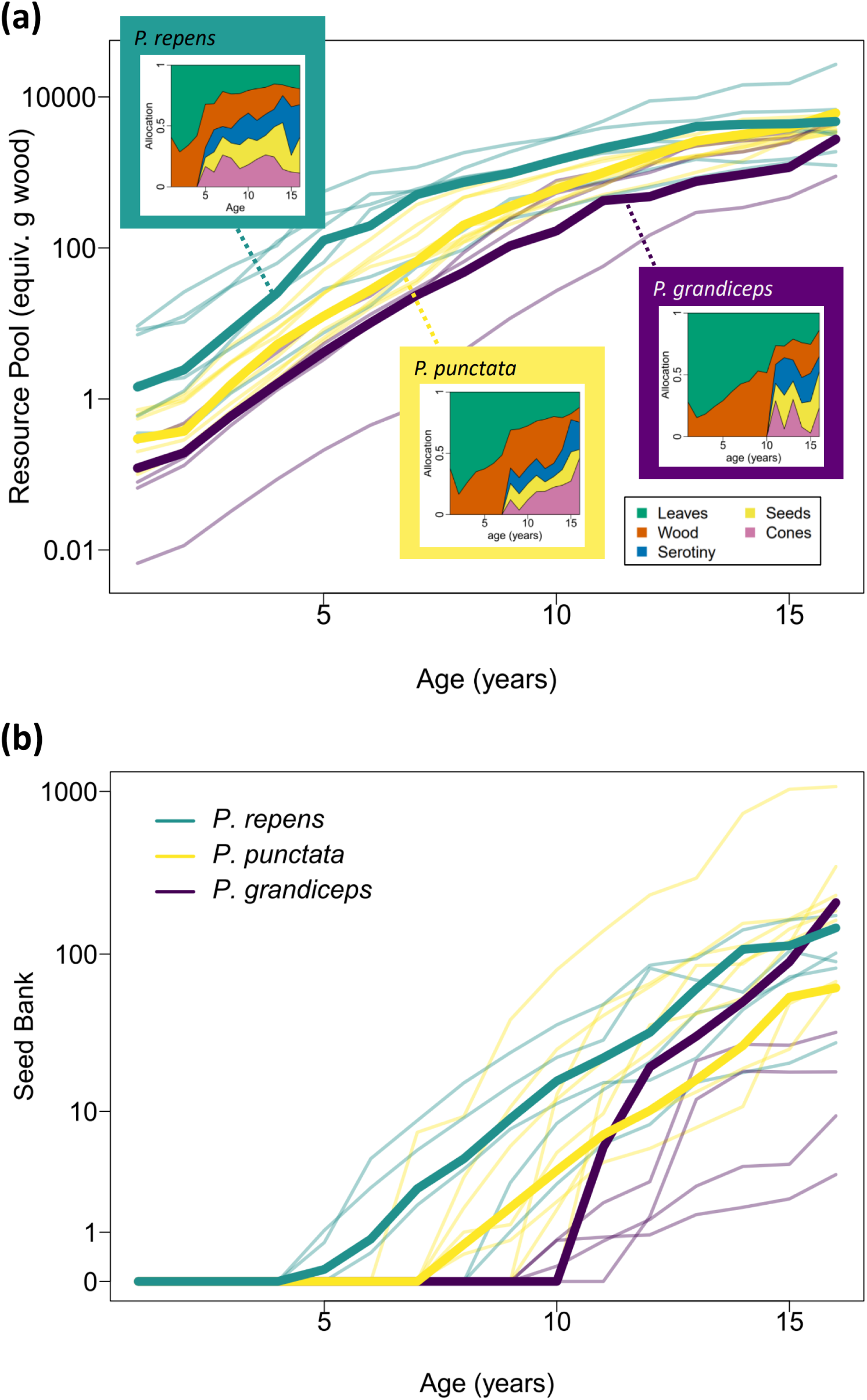
TRL model estimates of the age-dependent dynamics of whole-plant resource pools, resource allocation and total reproductive output (canopy seed bank size) for the 20 studied individuals of three *Protea* species on study site ‘jona_4’. (a) Posterior medians of resource pools versus age. Insets show resource allocation versus age for one individual per species. (b) Posterior medians of total reproductive output (size of the canopy seed bank) versus age for the same individuals.

### Trait effects on organ-level costs and benefits of life-history components

Functional traits affect organ-level costs and benefits of life-history components by-and-large in the same direction, such that trait values conferring greater benefits also increase costs (Fig. 7). Leaf construction cost increases with SLA, as does the rate of resource acquisition (Fig. 7a).

**Figure 7:**
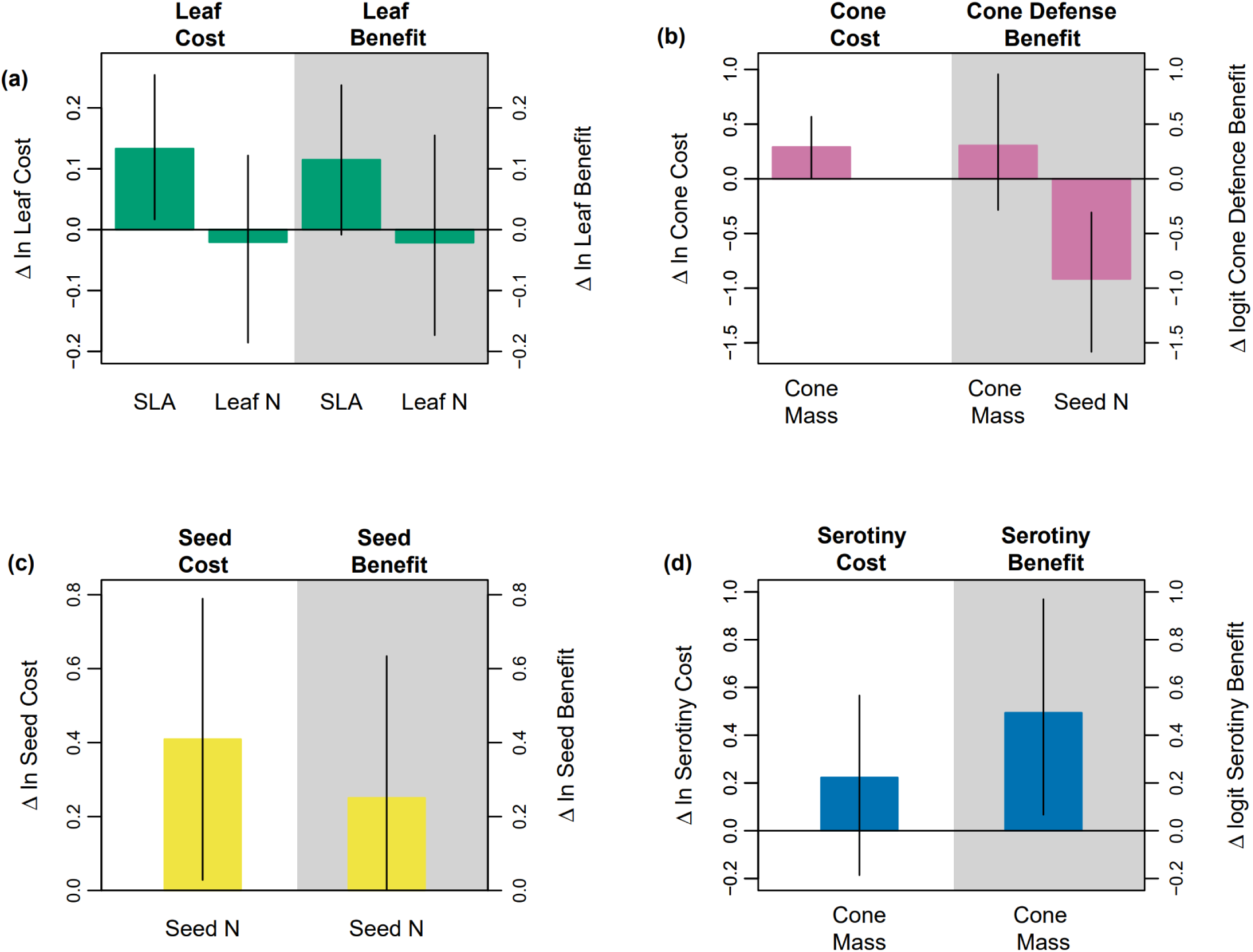
The effects of functional traits on paired organ-level costs and benefits of life-history components. Panels show posterior medians and 95% credible intervals of the effect of one standard deviation increase in ln trait values on the link-scale costs and benefits of (a) leaves, (b) cones, (c) seeds and (d) serotiny. Individual-level costs and benefits are estimated by the TRL framework applied to 22 species of *Protea*, inferred in an inverse-modelling approach from individual-level data on age-stratified performance measures and population-level trait values (except for leaf N concentration measured at species-level). For definitions of costs and benefits see ‘A Trait-Resource-Life-history (TRL) model for *Protea*’.

Heavier cones are more costly to produce and incur greater serotiny costs, but are more likely to remain closed and to be defended from seed predators (Fig. 7b,d). The greater cost of N-rich seeds is rewarded by the greater resource pool available to offspring (Fig. 7c) although this trades-off with an increased risk of seed predation (Fig. 7b).

### Relationships between organ-level costs and benefits, whole-plant resource budgets and whole-plant performance

Resource pools increase strongly with the amount of resources plants acquire from the seed, even well into the adult stage, and with the rate of resource acquisition from leaves (Fig. 8a). Resource pools also increase with leaf costs (since costly leaves are also more beneficial) and moderately with costs of cone production and serotiny (Fig. 8b). These patterns by-and-large hold with age, although the relationship between leaf cost and resource pools does tend to get stronger with time.

**Figure 8:**
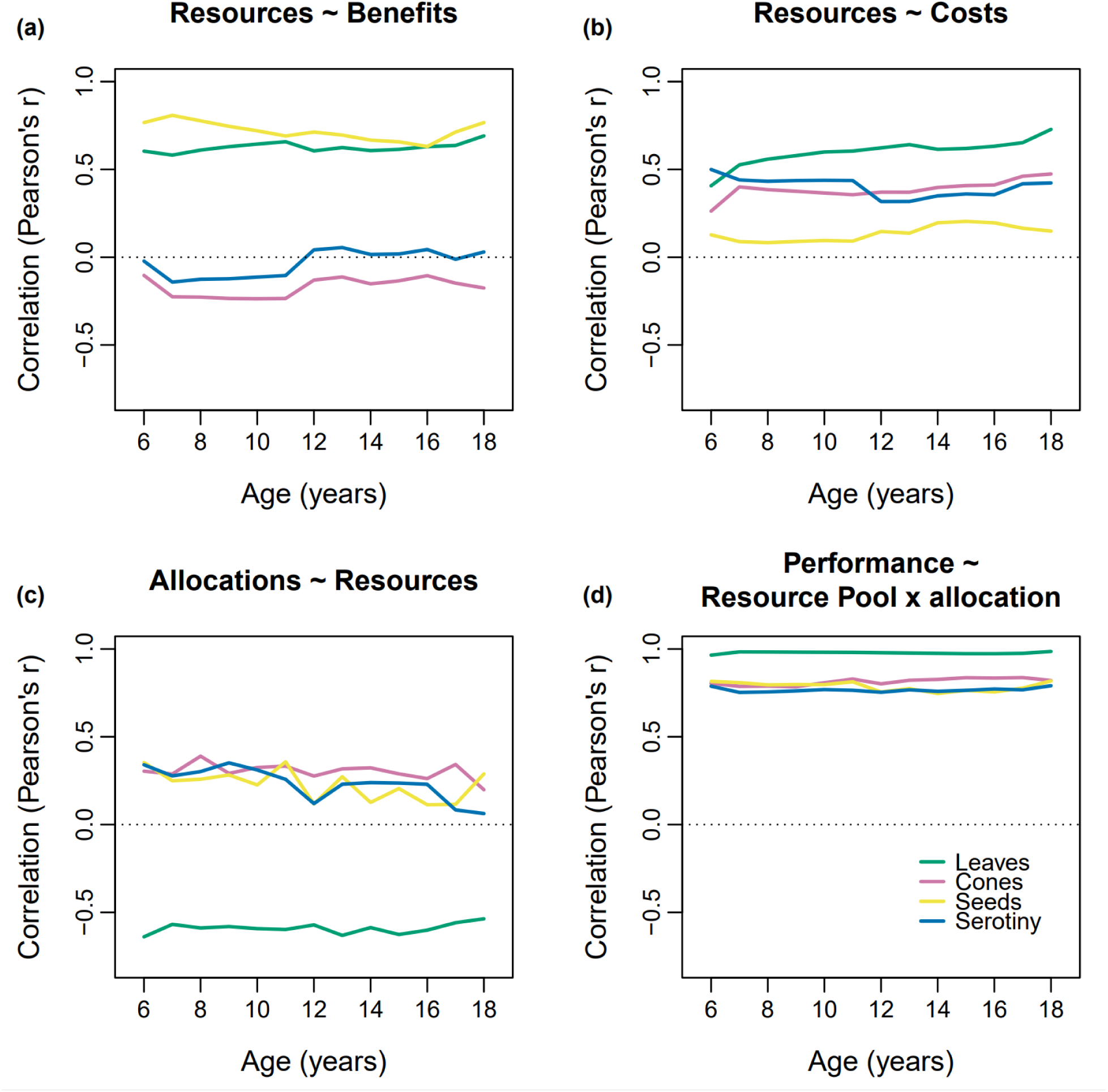
Age-specific relationships between organ-level costs and benefits, whole-plant resource budgets and whole-plant performance. Panels show Pearson’s correlation coefficients between (a) organ-level benefits and whole-plant resource pools, (b) organ-level costs and whole-plant resource pools, (c) whole-plant resource pools and allocations and (d) whole-plant resource investments (resource pool x allocations) and performance measures. Correlations were calculated from posterior median estimates for all 600 study individuals, using individual-level estimates of costs and benefits, and individual and age-specific estimates of resource pools, allocations and performance measures.

The larger the resource pool, the greater the share of resources plants allocate to reproduction and maternal care, and the lesser the share they allocate to production of leaves (Fig. 8c). These relationships only slightly weaken with age, at least for reproductive and maternal allocations. In turn, resource investment (the product of resource pools and resource allocation) is closely correlated with the respective life history component at all ages (Fig. 8d).

### Relationships of traits to whole-plant performance and life history

Leaf and seed benefits play a central role for resource budgets and performance at the whole plant level (Fig. 8). Hence, one may expect that SLA and seed N content, which shape these benefits (Fig. 7), should be correlated with whole-plant performance and other key life-history characteristics. SLA is indeed a reasonable predictor of all performance measures at early ages, but less so in later years, particularly for reproductive measures (Fig. 9a). Seed N content is, however, only weakly correlated with performance (Fig. 9b): leaf and seed production and serotiny decrease mildly with seed N, but wood and cone production are largely unrelated to seed N.

**Figure 9:**
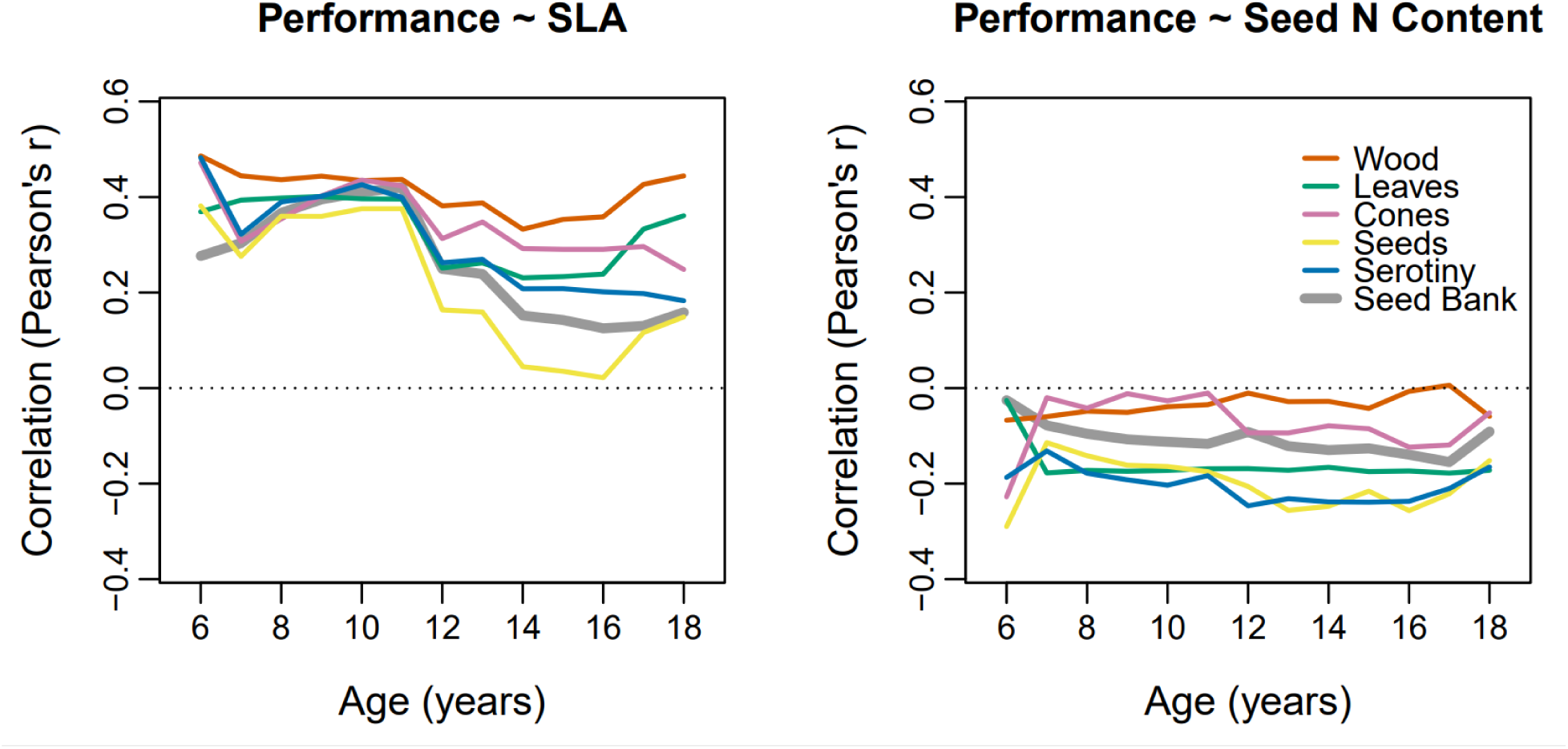
Age-dependent relationships between functional traits and whole-plant performance. Panels show Pearson’s correlation coefficients between whole-plant performance measures (including total reproductive output, an individual’ s seed bank) and (a) SLA and (b) seed N content. Correlations were calculated from posterior median estimates for all 600 study individuals, using individual-level estimates of whole-plant performance measures. Relationships between performance and the remaining functional traits are shown in Appendix S1: Figure S8).

Age of maturity as a key life-history characteristic shows clear relationships to a number of functional traits (Fig. 10;Appendix S1: Figure S6). In particular, it is negatively related to SLA across all study species and sites (Fig. 10a). Moreover, age of maturity is positively related with seed N content, although this relationship is weaker than for SLA (Fig. 10b). Traits shaping the costs and benefits of vegetative and reproductive organs thus play a substantial role for the pace of *Protea* life-histories. Moreover, correlations between traits and age of maturity have considerable conservation relevance since climate change is expected to increase fire frequency in much of the study region (Wilson *et al*., 2015). This should increase the extirpation risk of late-maturing populations (Enright *et al*., 2015) and lead to a functional shift of *Protea* communities towards species with higher SLA and lower seed N content. This is one example of how the TRL framework can contribute to a life-history based assessment of global change impacts on biodiversity (Ohse *et al*., 2023).

**Figure 10:**
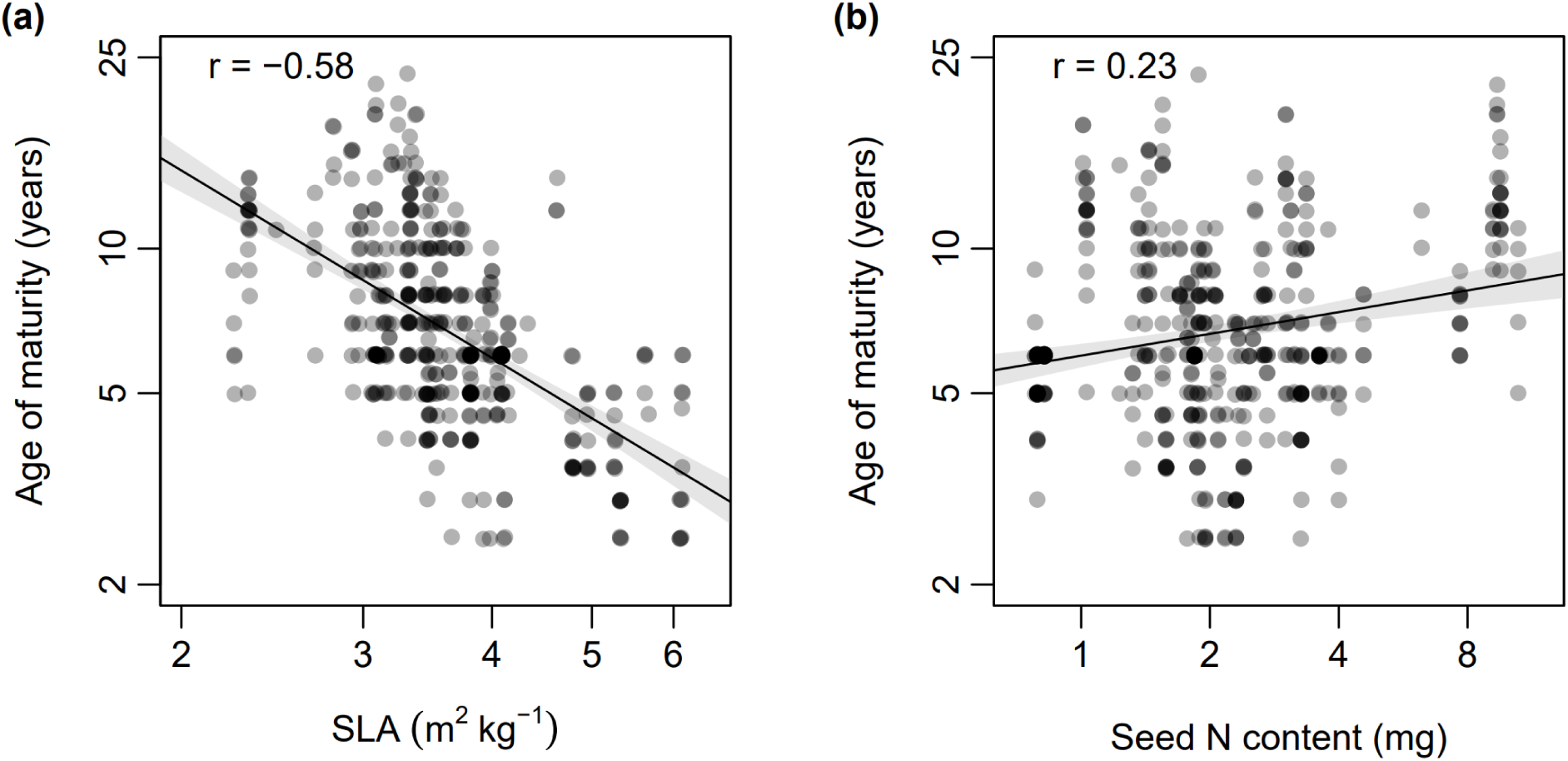
The relationship of (a) SLA and (b) seed N content with age of maturity for 600 individuals of 22 *Protea* species. The depicted age of maturity is the posterior median age at which an individual is predicted to have at least one seed in its canopy seed bank. Black lines show the fitted linear relationships between these variables and grey shadings show the 95% confidence intervals of these relationships. Pearson’s correlation coefficients between variables are shown in the top-left of each panel.

## Discussion

The TRL framework enables the formulation of integrated process-based models that predict the temporal dynamics of reproductive and vegetative performance throughout the lifetime of long lived plants. A TRL model describes how these temporal dynamics are shaped by the costs and benefits of plant organs, and by the dynamics of plant-level resource acquisition and allocation (Fig. 1). Importantly, the model can be parameterised with data on plant performance and functional traits (Fig. 4). This has three key benefits. First, we can predict performance measures with notable accuracy, throughout the lifetime of individuals up to the time of observation (Fig. 5). Second, by using a latent-variable modelling framework, we can learn about unmeasured variables that shape this process, namely, we can infer the resource costs and benefits of different plant organs (Fig. 7), the dynamics of resource budgets (Figs. 6,8), and unobserved whole plant performance measures such as the mass of leaves, the number of seeds produced annually, and total reproductive output (Fig. 8d,9). Third, by being able to infer these unmeasured variables, we can tease apart the processes linking organ-level traits to whole-plant-level performance and life-history. In the following sections, we discuss how the TRL model helps to infer the dynamics of whole-plant resource budgets and performance, how it contributes to a resource and trait based understanding of plant life-histories, how it can be applied to other plant systems and how it may promote a resource-based understanding of community dynamics.

### Inferring the dynamics of whole-plant resource budgets and performance

Understanding how functional traits shape life histories requires an understanding of how organ level traits scale-up to the lifetime dynamics of whole-plant resource acquisition, resource allocation and performance. By using hierarchical Bayesian latent variable modelling, the TRL framework can infer essentially unobservable variables (e.g. resource budgets), close gaps in performance observations for certain years (e.g. early growth and reproduction), and estimate performance quantities that are challenging to measure (e.g. the total mass of leaves produced per year). Further, by characterising all elements of a plant’s resource economy in consistent units (equivalents of woody biomass), with appropriate conversion parameters for other organs, we can directly compare the processes shaping life-histories in a single currency. With our case study, we have shown that the TRL framework can reconstruct the dynamics of resource budgets and performance from germination up to 33 years of age for the oldest individuals in the *Protea* dataset.

The construction costs of organs are often inferred from their mass and chemical composition at the time of harvest, using fixed conversion parameters to express costs in units of glucose (Poorter, 1994; Williams *et al*., 1997). The TRL framework takes an inverse approach, estimating net costs and benefits over the lifetime of an organ, based on the effect of organs on whole plant resource pools. This inherently accounts for the costs of organ maintenance, and the benefits of nutrient recycling (e.g. from senescing leaves) that are not considered in once-off construction cost calculations (Poorter, 1994). The inverse approach taken by TRL also accounts for the different costs of nutrient acquisition experienced by different species in different environments, a prohibitive task with fixed conversion parameters (Poorter, 1994). Further, within this integrated framework one can estimate the benefits of leaf production for resource acquisition, of seed resources for seedling growth, and of resource allocation to serotiny and to defence against seed predation. While the first two of these could be estimated via extensive physiological measurements, the presented integrative approach enables us to estimate these benefits from data that are easier to obtain.

### Resource and trait-based understanding of plant life-histories

A resource-based understanding of life histories is facilitated by the TRL framework’s description of how organ-level costs and benefits shape whole-plant resource budgets and performance (Fig. 1). Applying this framework to *Protea*, we found that higher resource acquisition be it initial acquisition from the seed or subsequent acquisition through leaves is linked to the build-up of larger resource pools at all ages (Fig. 8a). The size of the resource pool is in turn linked to vegetative versus reproductive allocation: plants with more resources invest a larger share of these resources into reproduction (Fig. 8c). Since performance is strongly correlated with resource investment (the product of resource pool and resource allocation; Fig. 8d), a greater resource pool implies a disproportionate increase in reproductive performance.

Functional traits are relatively modest predictors of whole-plant performance (Fig. 9), even if they play important roles for the organ-level costs and benefits that shape resource budgets and life histories (Fig. 7). This is because multiple co-varying processes are involved in upscaling from organ-level traits to whole plant resource budgets, which themselves vary through time and are allocated to multiple dimensions of performance. For example, seed N content increases seed benefits that are linked to greater whole-plant resource pools and performance, but it also increases the costs of both seeds and cones (Figs. 7,8). Overall, this leads to negative and relatively weak relationships between seed N content and all performance measures (Fig. 9b). On the other hand, SLA as the key trait determining resource acquisition from leaves (Fig. 7), is correlated with earlier maturity (Fig. 10a) and showed strongly positive relationships to all performance measures at young to intermediate ages (Fig. 9a). However, the positive relationship between SLA and total reproductive output clearly weakened for older ages (Fig. 9a). Hence, traits do matter (Adler *et al*., 2013), but in ways that cannot necessarily be captured by simple correlations with whole-plant performance (Kleyer & Minden, 2015; Messier *et al*., 2017; Swenson *et al*., 2020).

The TRL framework should enable the detection of trade-offs at the level of organs, whole-plant resource budgets and life histories. At organ level, we found clear trait-mediated trade-offs: functional trait effects on the costs and benefits of an organ generally act in the same direction (Fig. 7). Hence, if a plant is to gain the benefits of an organ, it has to invest resources to reap this reward. This is a fundamental tenet of evolutionary life-history theory (Stearns, 1992) – resource investment into an organ that is not sufficiently beneficial to fitness will be selected against.

When examining resource budgets, we also detected a clear trade-off between resource allocation to growth versus reproduction (Fig. 8c). Such a growth-reproduction trade-off may give rise to a life-history-level trade-off between early and late reproduction (Stearns, 1992; Salguero-Gómez *et al*., 2016). This is expected if the earlier maturity of plants that start with many resources reduces their subsequent resource acquisition and reproduction (Tonnabel *et al*., 2012). In our case study, we found no evidence for this life-history trade-off: plants that matured earlier generally also reproduced more later in life (Appendix S1: Figure S7). This may be because plants performing well throughout their lives have continuously higher resource acquisition (van Noordwijk & de Jong, 1986) that results from factors not considered in our analysis, e.g. because they occupy more resource-rich microsites or experience less competition (Walter *et al*., 2023). A comprehensive test for the trade-off between early and late reproduction should thus aim to standardize resource availability and/or account for the competitive environment of plant individuals. While we did not detect a trade-off between early and late reproduction in the *Protea* system, there is evidence for a life-history trade-off between the quantity and quality of offspring: plants with high seed N content mature later and produce fewer seeds (Fig. 9b, 10b), but their seedlings have a higher survival probability (Cooksley *et al*., 2023).

### Application of the TRL framework to other plant systems

Application of the TRL framework to other study systems will help to further develop the framework and extend its general validity, advancing a general understanding of how functional traits shape resource dynamics and performance of plants. This transfer to other study systems requires data on functional traits as proxies for costs and benefits of resource investment into organs, data on the temporal dynamics of life-history components and – potentially – data on the temporal dynamics of resource budgets (Fig. 1). It seems worth noting that the parameterisation of a TRL model using latent state-space modelling does not require a comprehensive coverage of all of these data types at all times.

TRL considers functional traits as measures for the costs and benefits of investment into different life-history components (Fig. 1). These life-history components are typically linked to a particular organ type (such as cones, seeds, wood or leaves). Biomass or nutrient investment at organ-level can be captured by a few well-established traits such as SLA, wood density or seed (nutrient) mass. Hence, application of TRL models should typically require a limited number of traits for which measurements are widely available (Kattge *et al*., 2020) or can be taken relatively easily. Moreover, since a given trait often affects both the costs and the benefits of an organ (Fig. 7; see also Westoby *et al*., 2002; Wright *et al*., 2004; Chave *et al*., 2009), application of the TRL framework should typically require easily accessible data on a limited number of traits.

Reconstruction of vegetative growth in *Protea* individuals relied on their regular growth pattern (Fig. 3). Other species-rich genera in the Proteaceae show a similar growth pattern so that our approach should be easily transferable to them (e.g. Australian *Banksia* and South African *Leucadendron*; Lamont, 1985; Harris & Pannell, 2010). Beyond the specific case of Proteaceae, vegetative growth of a broad range of tree species can be reconstructed by combining dendrochronological data with allometric relationships between trunk diameter and wood or leaf biomass (Babst *et al*., 2016). A similar approach should be feasible for herbaceous perennials from seasonal environments that commonly form annual growth rings in the main root (Dietz & Ullmann, 1997).

For long-lived plants, data on the temporal dynamics of reproduction are generally more difficult to obtain than data on vegetative growth. In many serotinous plants, however, past reproduction and maternal care can be quantified relatively easily. Serotiny is no exception in the plant kingdom: it occurs in more than 40 genera that play important roles in both Mediterranean-type ecosystems and coniferous forests (Lamont *et al*., 1991; 2019). It thus seems both interesting and relevant to apply the TRL framework to a broader range of serotinous species.

Non-serotinous trees arguably pose one of the biggest challenges for reconstructing lifetime reproductive schedules. However, long-term records of individual reproductive output exist for certain tree species (Bogdziewicz *et al*., 2020). In other species, it is possible to reconstruct components of past reproduction. For instance, annual cone production in pinyon pines (*Pinus edilis*) can be reconstructed from cone abscission scars (Redmond *et al*., 2016). Moreover, remote sensing can be used to estimate current reproductive output. For instance, terrestrial lidar scanning of a *Pinus pinea* stand quantified canopy characteristics that were informative of between-individual variation in cone number and cone mass (Schneider *et al*., 2020). Remote sensing methods can thus yield snapshots of reproductive output for large numbers of individuals. Such data may be used to infer reproductive schedules at population or species level using individual-for-age substitution. Alternatively, the reproductive output of trees in a given year can be estimated inversely by combining data on spatial distribution of adult trees and post dispersal seed densities (e.g. Clark *et al*., 2019). Such estimates can be further improved by marker-based genotyping of seeds and adults and by integrating dendrochronological data for adults (Lamonica *et al*., 2021).

In short-lived plants, it is possible to directly observe the temporal dynamics of reproduction and vegetative growth. Notably, rapid developments in noninvasive plant phenotyping (Pieruschka & Schurr, 2019) make it possible to efficiently observe the temporal dynamics of multiple life history components at high temporal resolution (e.g. Vasseur *et al*., 2019). Short-lived plants also offer the possibility of experimentally controlling resource availability for individuals to rigorously test for life-history trade-offs (see *Resource and trait-based understanding of plant life-histories*). This is particularly true for the model species *Arabidopsis thaliana*, for which it is even possible to manipulate both resource acquisition and allocation (Schulze *et al*., 1994; Bennett *et al*., 2012). This opens interesting avenues for experimentally testing the TRL framework.

### Resource-based understanding of community dynamics

A key aim of trait-based ecology is to use functional traits to explain the structure and dynamics of higher levels of biological organisation (Lavorel & Garnier, 2002; McGill *et al.,* 2006; Salguero-Gómez *et al*., 2018). This requires scaling-up through the levels of this hierarchy, in which emergent properties at higher levels are shaped by the integration of, and covariances between, multiple lower-level processes (Laughlin & Messier, 2015; Shipley *et al.,* 2016). By combining the TRL framework, which describes the scaling-up from organs to individual-level resource budgets, with descriptions of resource-based interactions among individuals (Mittelbach & McGill, 2019), it should be possible to develop integrative, resource-driven models of community dynamics.

In our model system, explicitly considering resource flows in analyses of demography and community assembly has proven fruitful. Traits shaping competition for abiotic resources determine interactions and performance at both the adult (Walter *et al*., 2023) and juvenile (Cooksley *et al*., 2023) stage of *Protea* individuals. Furthermore, floral and seed resources at plant-, local and site-levels shape the interaction frequencies of *Protea* individuals with animal mutualists and antagonists (Schmid *et al*., 2015; Nottebrock *et al*., 2017; Neu *et al*., 2023a,b) as do the energetic requirements of the interacting animals (Neu *et al*., 2023a). In turn, the foraging of animal mutualists and antagonists affects *Protea* fitness via pollination and seed predation, respectively (Nottebrock *et al*., 2017; Walter *et al*., 2023).

An ambitious, but logical, next step would be to integrate these disparate analyses via a latent variable modelling framework, driven by the flow of resources from the abiotic environment over competing plants to higher trophic levels. This could be done by integrating resource-driven interactions into the TRL framework to scale up resource dynamics from organisms to entire communities. Modelling such hierarchical dynamics via a universal currency of resources would have clear advantages, particularly avoiding the intractability of quantifying pairwise interaction coefficients among species in diverse communities (Broekman *et al*., 2019). Ultimately, such an integrative resource-driven understanding of life-histories and community dynamics may feed into a new generation of ecosystem models that represent biodiversity by general ecological mechanisms rather than a predefined set of coarse functional strategies of organisms (Scheiter *et al*. 2013; Harfoot *et al*., 2014; Higgins, 2017).

## Conclusions

The proposed TRL framework enables data-driven inference on the processes linking traits, resource budgets and life-histories of long-lived plants. Its application to a broad range of study systems and its integration with models of community dynamics hold promise for a process based understanding of higher levels of biological organisation and a closer theory-founded integration of functional ecology, evolutionary ecology, community ecology and ecosystem science.

## Acknowledgements

Fieldwork was conducted under Western Cape Nature Conservation Board permit 0028- AAA008-00262 and the following landowners and reserves gave permission to work on their properties: Ceres Mountain Fynbos Nature Reserve and Witzenberg Municipality, City of Cape Town and Helderberg Nature Reserve, Fernkloof Nature Reserve and Overberg Municipality, Flower Valley Conservation Trust, Grootbos Private Nature Reserve, Heuningklip Farm, Hottentots Holland Nature Reserve, Limietberg Nature Reserve, Mont Rochelle Nature Reserve and Riviersonderend Nature Reserve. Sarah de Gruchy, Madelein de Klerk, Nicolene Hellström, Liezel Knight, Michael Leach, Marisa Noordergraaf, Hannah Oliphant, Barbara Seele, Adrian D.A. Simmers, Megan Smith and Martina Treurnicht provided invaluable help with field and lab work. We thank the Central Analytical Facility of the University of Stellenbosch for nutrient determination, and Carsten Buchmann, Jörn Pagel and Hanna Walter for helpful discussions.

This research was funded by the German Research Foundation (DFG) project ProteaNet (grant numbers SCHU 2259/3-3 and SCHL 1934/1-3).

## Author contributions

Huw Cooksley, Matthias Schleuning and Frank Schurr designed the study and the conceptual TRL framework. Huw Cooksley and Alex Neu collected data for the *Protea* case study, and Huw Cooksley and Frank Schurr performed the analysis. Huw Cooksley and Frank Schurr led the writing of the manuscript, and all authors contributed revisions and accepted the final version.

## Open research statement

Data as well as JAGS and R code for fitting the TRL model for the *Protea* case study are provided in the supplementary material accompanying this article.

