## Appendix S1 supplementary figures and tables for "An integrative trait-based framework to infer resource budgets and life-histories of long-lived plants"

**Appendix S1, supplementary figures and table for:**

### An integrative trait-based framework to infer resource budgets and life-histories of long-lived plants

Huw Cooksley[1,2], Matthias Schleuning[3], Alexander Neu[2,3,4], Karen J. Esler[2], Frank M. Schurr[1,5]

1: Institute of Landscape and Plant Ecology, University of Hohenheim, Stuttgart, Germany

2: Department of Conservation Ecology and Entomology, Stellenbosch University, Stellenbosch, South Africa

3: Senckenberg Biodiversity and Climate Research Centre (SBiK.F), Frankfurt am Main, Germany

4: Institute for Ecology, Evolution and Diversity, Goethe University Frankfurt, Frankfurt am Main, Germany

5: KomBioTa – Center for Biodiversity and Integrative Taxonomy, University of Hohenheim & State Museum of Natural History, Stuttgart, Germany

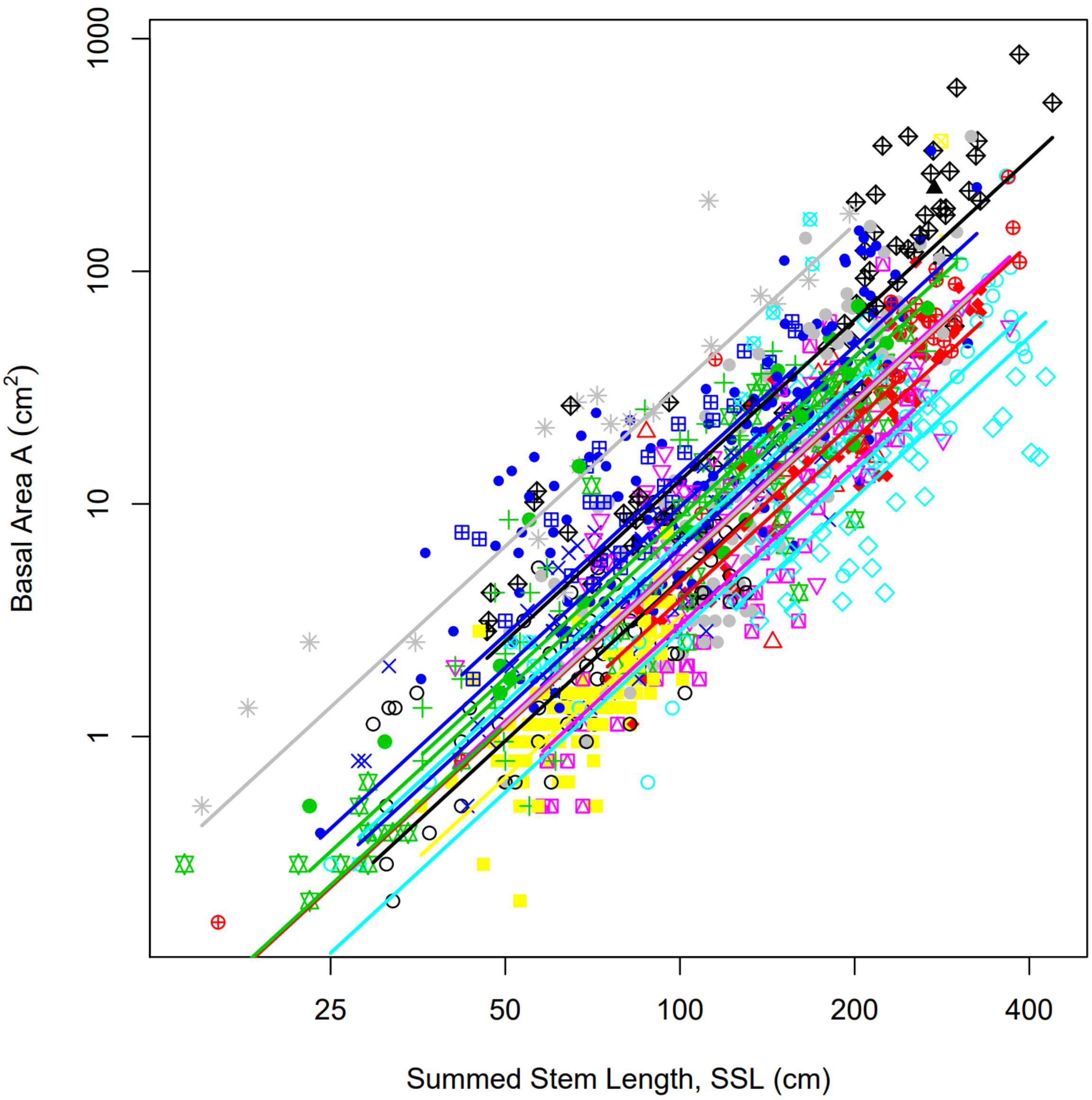


Figure S1: Allometric relationships between stem basal area (A) and summed stem length (SSL) for 1,041 individuals across the 22 species used in the *Protea* case study. Lines show species-specific fitted relationships from a mixed standardised major axis regression between A and SSL, with random intercepts for species. Points of differing shape and colour represent the 22 species, and lines are coloured by species with the same colour scheme.

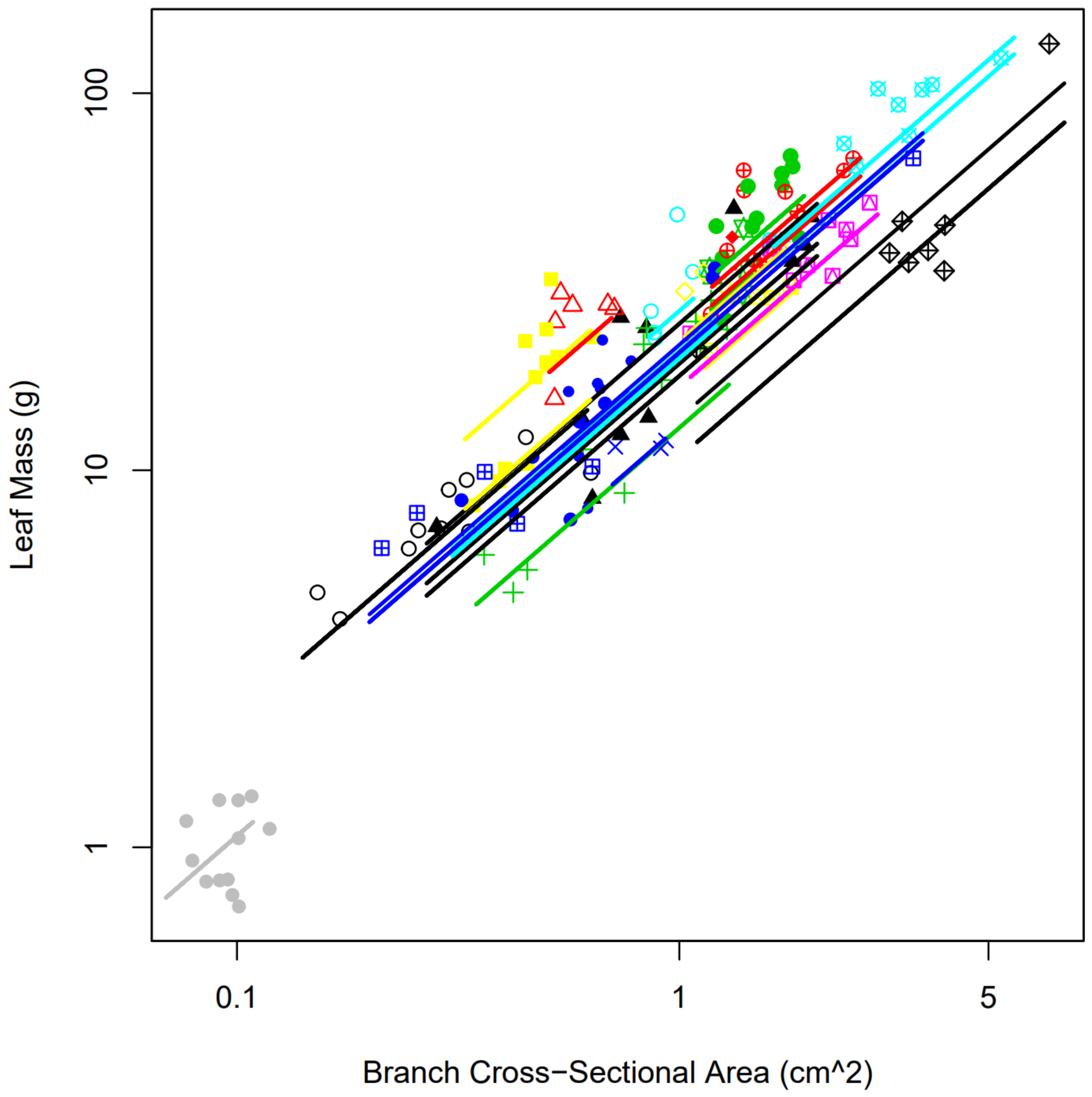


Figure S2: Allometric relationships between stem leaf mass per branch and branch cross-sectional area, across the 22 species used in the *Protea* case study. Lines show population-specific fitted relationships from a mixed standardised major axis regression between leaf mass and cross-sectional area, with random intercepts for species and site, from measures pooled for individual. Points of differing shape and colour represent the 22 species, and lines are coloured by species with the same colour scheme.

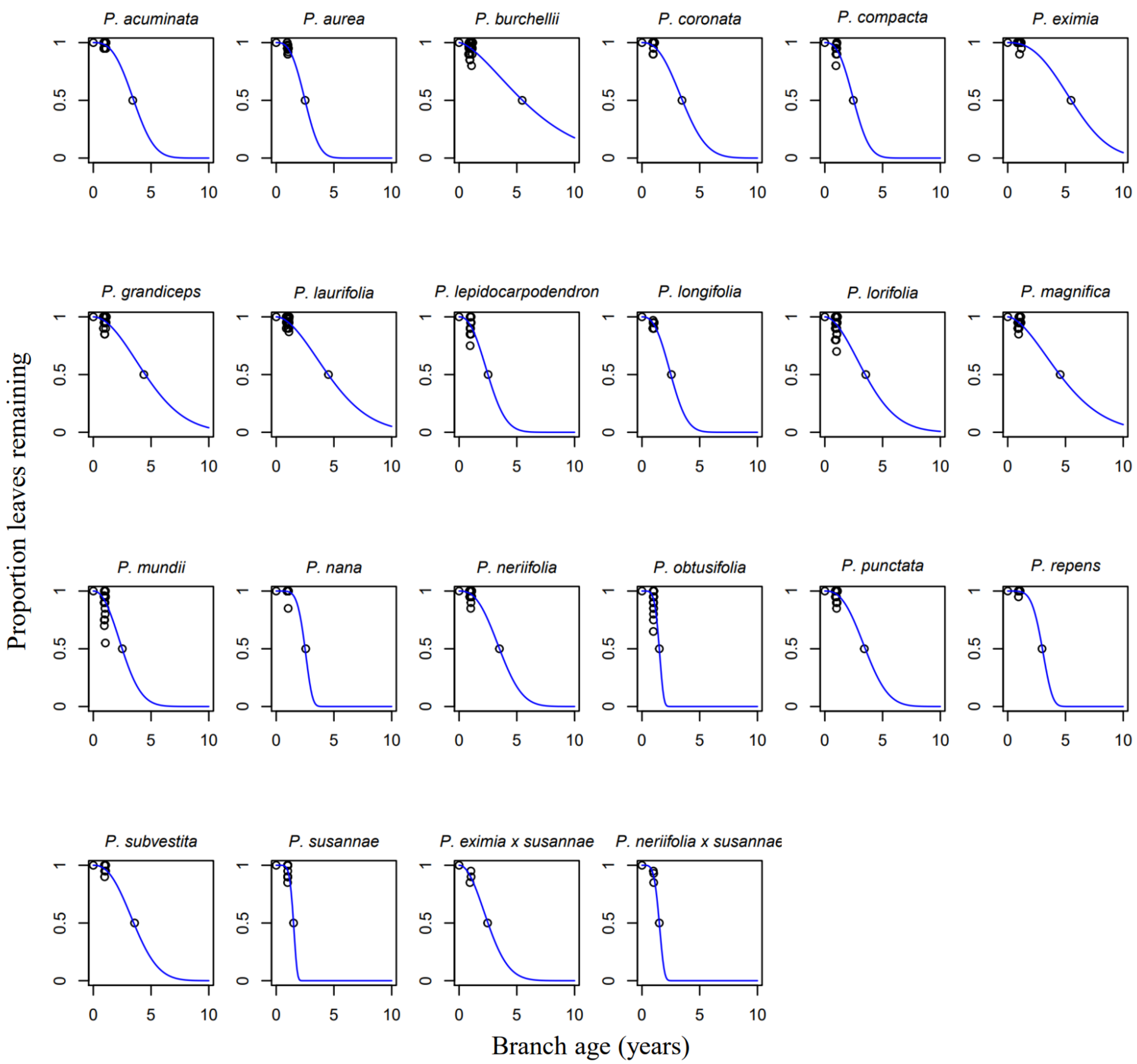


Figure S3: Leaf retention curves for all 22 species used in the *Protea* case study. Each panel shows the estimates of proportion of produced leaves remaining on a branch with branch age for each species, with curves representing the species-specific non-linear least squares fits of the complement of the Weibull's cumulative density function to this data.

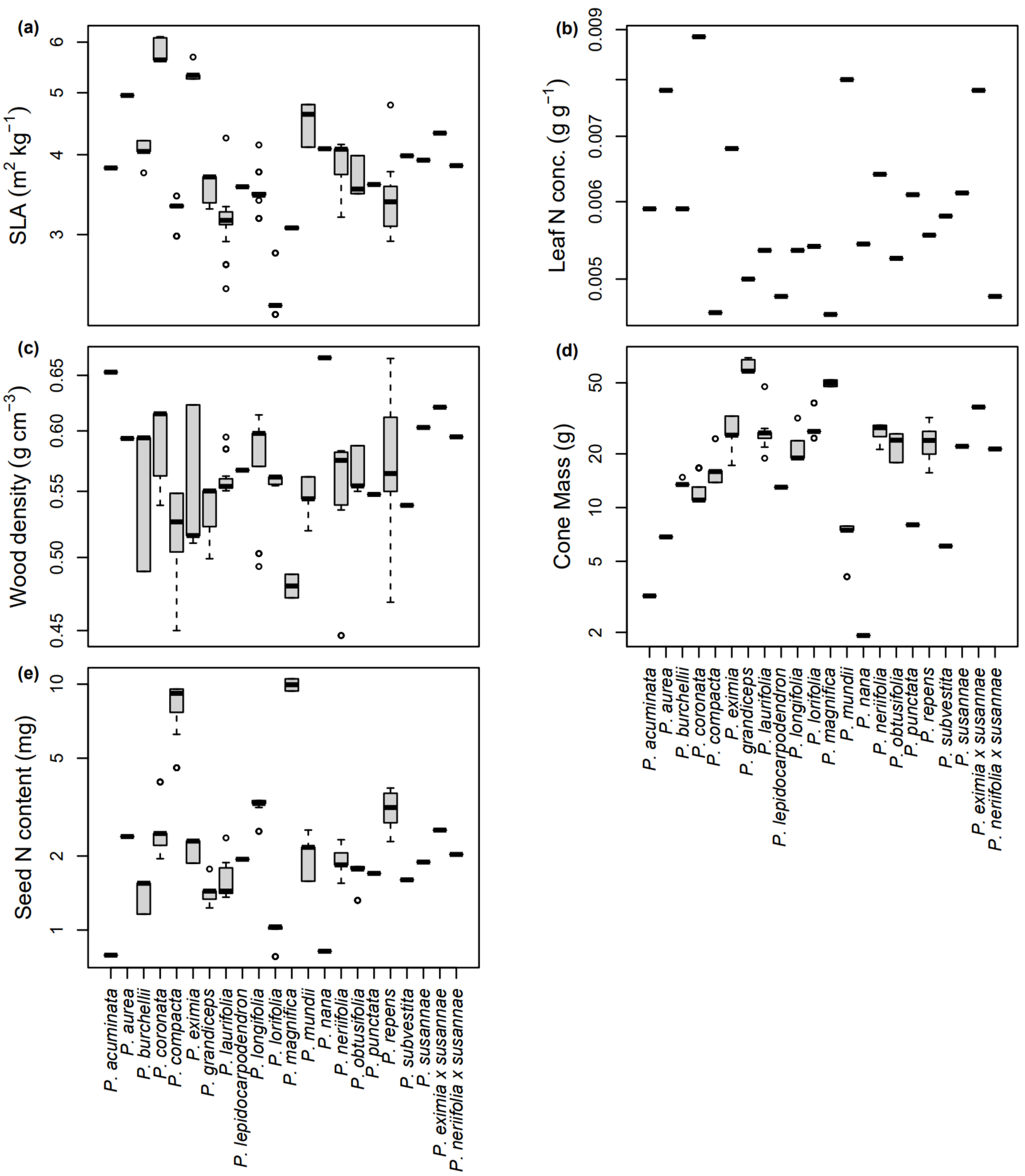


Figure S4: Variation within and between species for the functional trait values used in the *Protea* case study, for the traits (a) SLA, (b) leaf N concentration, (c) wood density, (d) cone mass and (e) seed N content. We used species- and site- mean trait values for all traits except leaf N concentration (species-mean). Bold lines show medians, boxes the interquartile range, whiskers extend this range up to 1.5-fold on either side, and dots are outliers.

| Functional trait | N measures | N sites |
| --- | --- | --- |
| Wood density | 389 | 21 |
| SLA | 897 | 21 |
| Leaf N concentration | 192 | 28 |
| Cone mass | 517 | 21 |
| Seed N content | 77 * | 21 |

Table S1: Sample sizes for functional traits used in the *Protea* case study. N measures refers to the number of individuals on which a trait was measured. For seed N content determination, seed samples were pooled by population, thus N measures refers to the number of populations sampled for this trait.

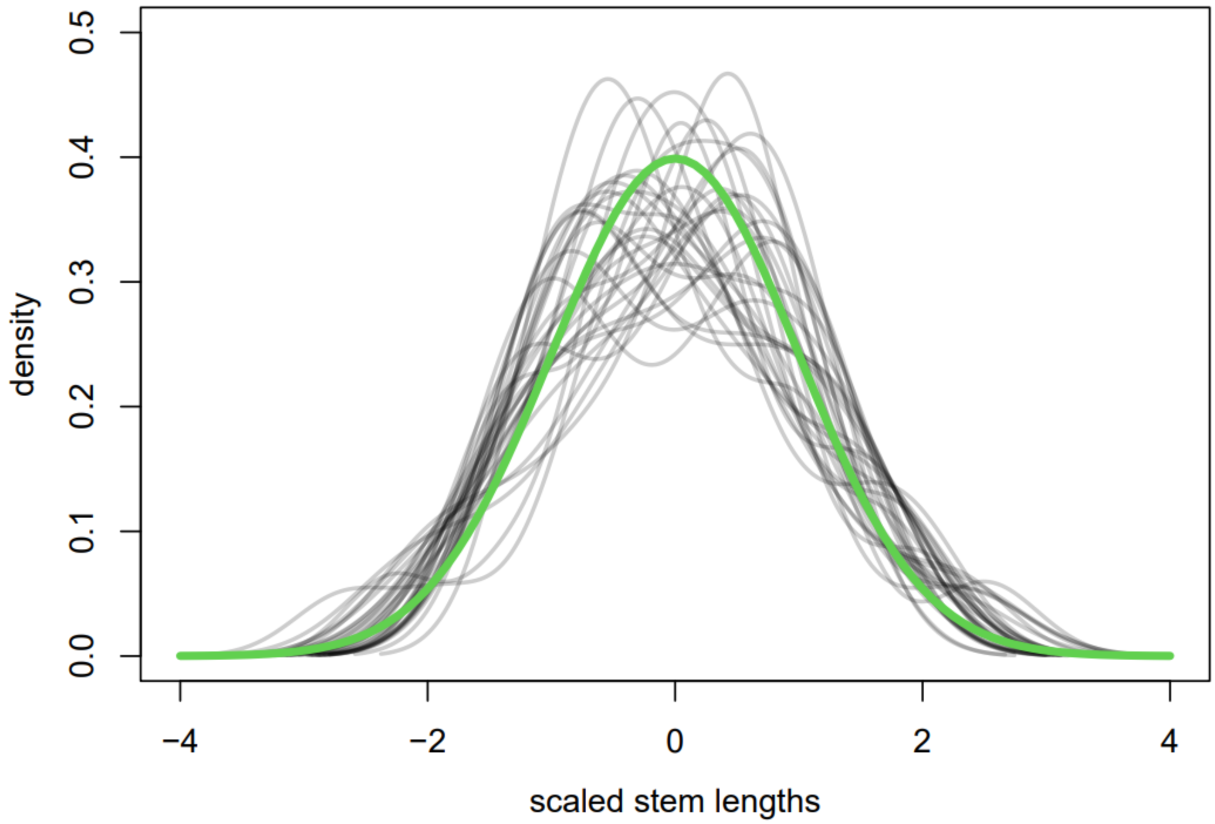


Figure S5: Length of stem increments across a year within a plant approximately follow a normal distribution with variance equal to the mean. Each black line shows the scaled (mean=0, sd=1) distribution of stem increments produced in the same year on a plant, measured from the study species of the *Protea* case study. The green line shows the standard normal distribution. For clarity, we show 30 distributions only.

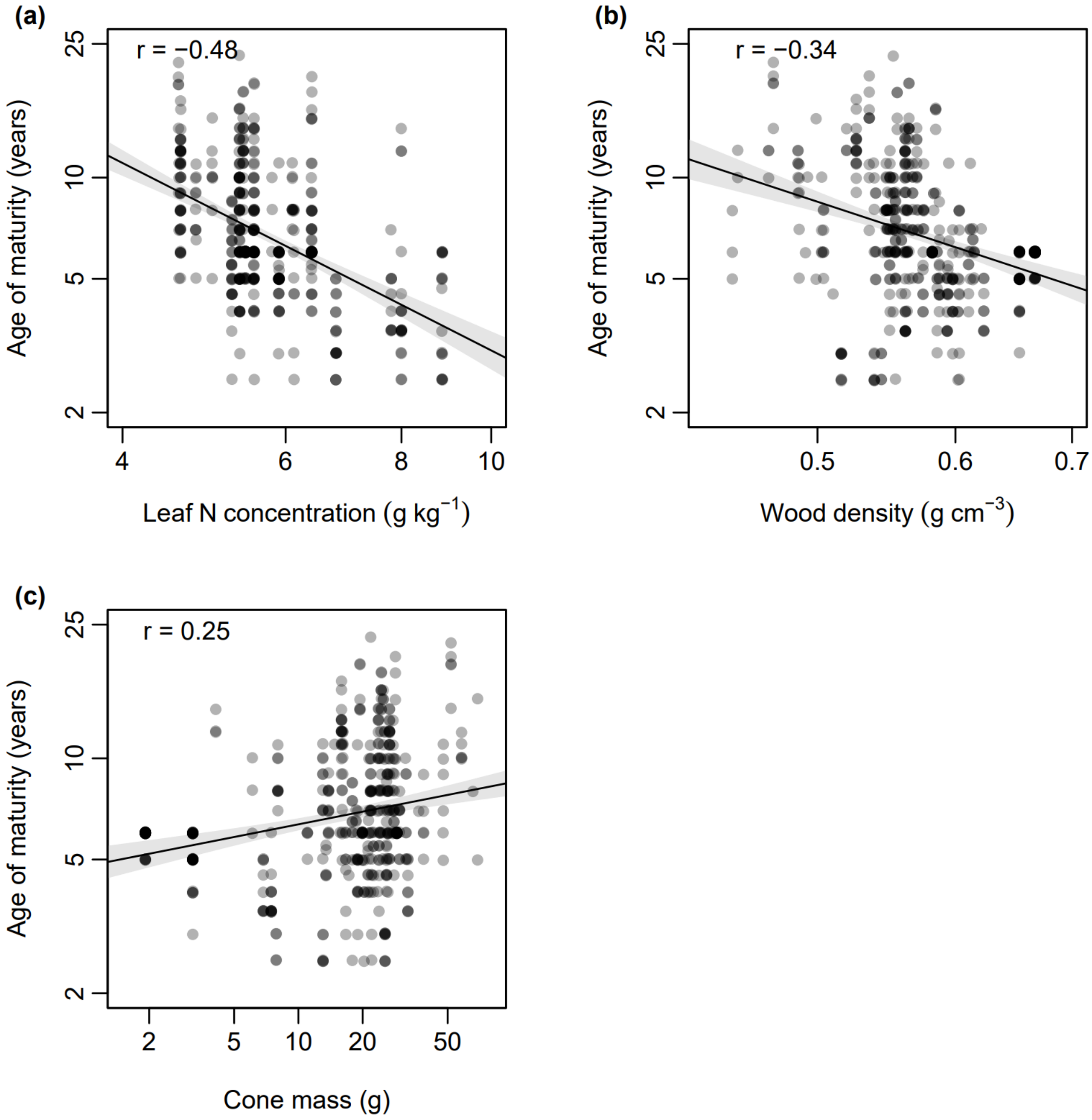


Figure S6: The relationship of (a) leaf N concentration, (b) wood density and (c) cone mass with age of maturity for 600 individuals of 22 *Protea* species. The depicted age of maturity is the posterior median age at which an individual is predicted to have at least one seed in its canopy seed bank. Black lines show the fitted linear relationships between these variables and grey shadings show the 95% confidence intervals of these relationships. Pearson's correlation coefficients between variables are shown in the top-left of each panel.

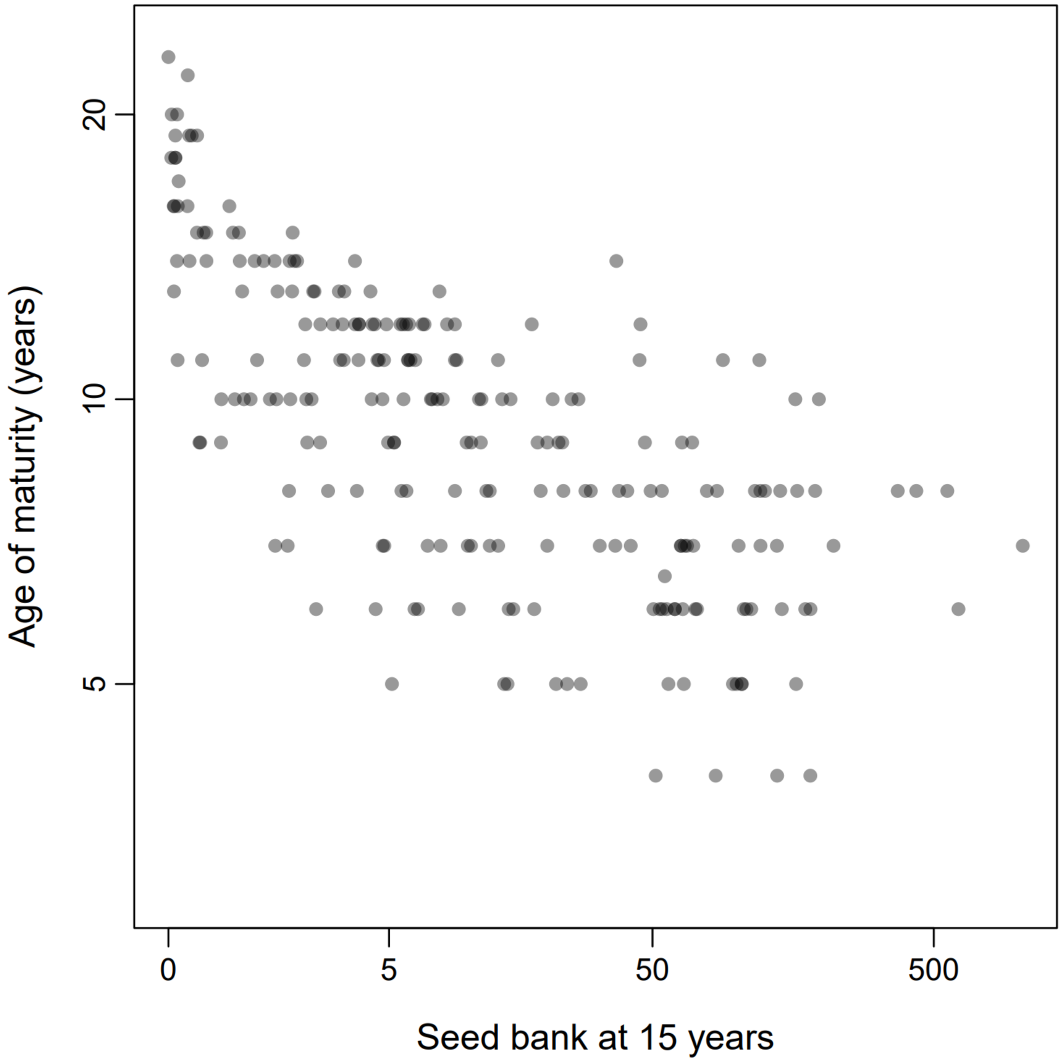


Figure S7: The relationship of age of maturity with seed bank at 15 years, for 600 individuals of 22 *Protea* species. The depicted age of maturity is the posterior median age at which an individual is predicted to have at least one seed in its canopy seed bank

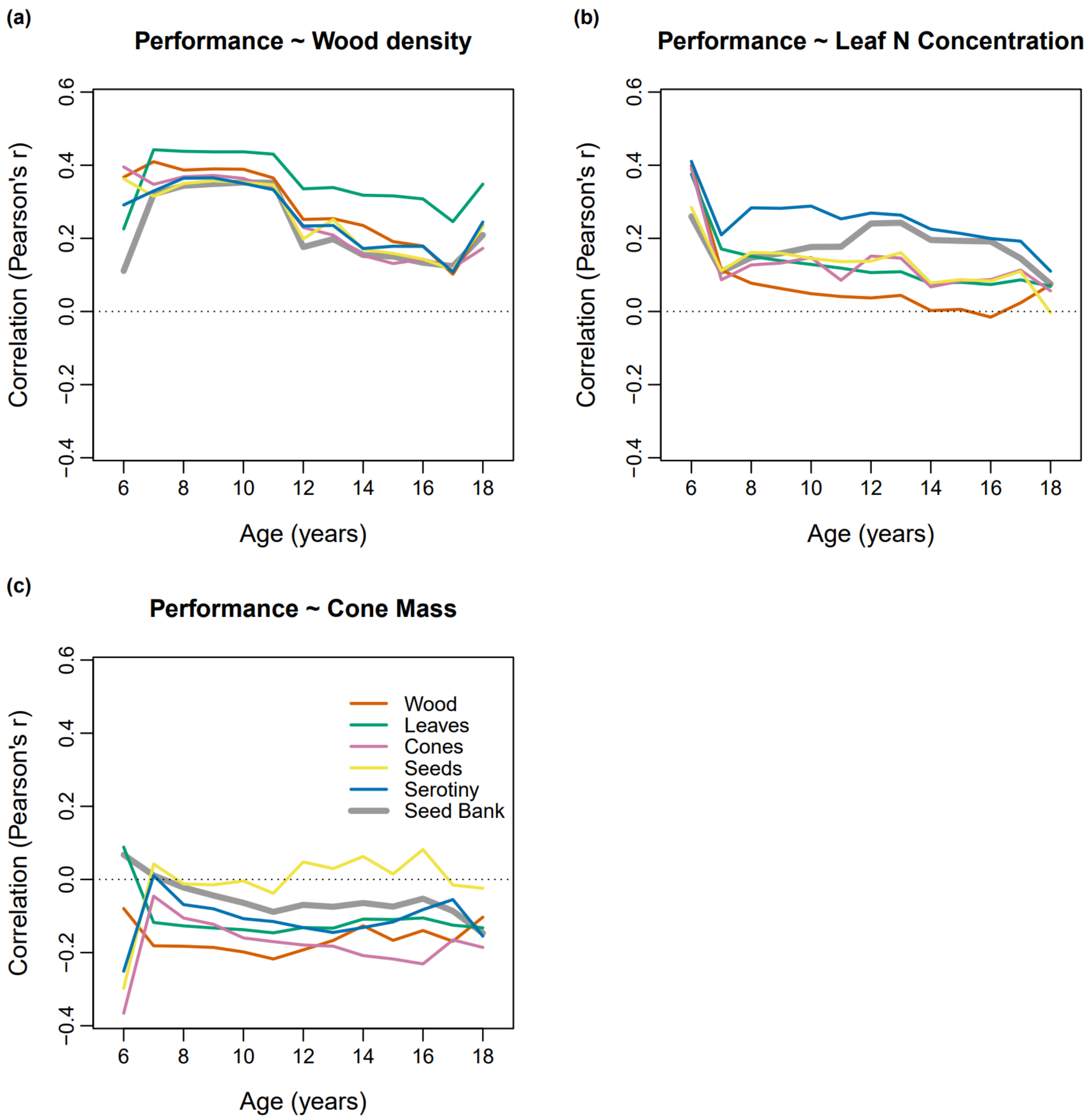


Figure S8: Age-dependent relationships between functional traits and whole-plant performance. Panels show Pearson's correlation coefficients between whole-plant performance measures (including total reproductive output, an individual's seed bank) and (a) wood density, (b) leaf N concentration and (c) cone mass. Correlations were calculated from posterior median estimates for all 600 study individuals, using individual-level estimates of whole-plant performance measures.
