## Appendix S2 supplementary code for "An integrative trait-based framework to infer resource budgets and life-histories of long-lived plants"

### Appendix S2, supplementary code to fit the TRL Protea case study, for:
###
###  'An integrative trait-based framework to infer resource budgets and life-histories of long-lived plants'
###
###   Huw Cooksley1,2, Matthias Schleuning3, Alexander Neu2,3,4, Karen J. Esler2, Frank M. Schurr1,5
###   
###   1: Institute of Landscape and Plant Ecology, University of Hohenheim, Stuttgart, Germany
###   2: Department of Conservation Ecology and Entomology, Stellenbosch University, Stellenbosch, South Africa
###   3: Senckenberg Biodiversity and Climate Research Centre (SBiK.F), Frankfurt am Main, Germany
###   4: Institute for Ecology, Evolution and Diversity, Goethe University Frankfurt, Frankfurt am Main, Germany
###   5: KomBioTa - Center for Biodiversity and Integrative Taxonomy, University of Hohenheim & State Museum of Natural History, Stuttgart, Germany


# setwd("...") # please change as appropriate

library(saveJAGS)

load("TRL_data.rdata")


### Specify JAGS code for the TRL model Protea case study
cat("
model {
  
  # Loop through individuals
  for (i in 1:NInds){
    
    # Need to set SSL variables in thee first time-step to zero.
    # For these variables we need to call the value in the previous time-step.
    # Thus t=1 really starts in the second row for these variables.
    SSL[1,i] ~ dbin(0,0)
    SSLobs[1,i] ~ dbin(0,0)


    ##********************##
    ## Costs and benefits ##
    ##********************##
    
    ## Seed Benefit and resource pool in the first time-step
    logR[1,i] ~ dnorm(mu_r1[i], tau_r1)
    mu_r1[i] <-
      alpha_sb +
      alpha_sb_sp[Species[i]] +
      alpha_sb_site[Sites[i]] +
      d1 * zeros_vec[i] +
      (beta_snmsb + beta_snmsb_si[Sites[i]]) * scc_log_SeedNM[i] +
      exp(log_b_pa + log_b_pa_sp[Species[i]] + log_b_pa_si[Sites[i]]) * delta_t[i]

    # NB: the 'd1 * zeros_vec' term (and the corresponding term in other costs/benefits) is a handy trick
    # to get JAGS to block-update the cost and benefit parameters using the glm sampler, allowing 
    # efficient mixing. The likelihood is unafected, as this term is equal to zero


    logSeedBenefit[i] <- logR[1,i] -
      exp(log_b_pa + log_b_pa_sp[Species[i]] + log_b_pa_si[Sites[i]]) * delta_t[i]
      


    ## Leaf Cost
    logLeafCost[i] ~ dnorm(
      alpha_lc +
      alpha_lc_sp[Species[i]] +
      alpha_lc_si[Sites[i]] +
      d2 * zeros_vec[i] +
      beta_csla * scc_log_SLA[i] +        # the 'scc_log_' prefix simply means log-tranformed, then scaled and centred
      beta_cln * scc_log_LeafN[i],
      tau_lc)


    ## Leaf Benefit
    logLeafBenefit[i] <- 
      alpha_lb +
      alpha_lb_sp[Species[i]] +
      alpha_lb_si[Sites[i]] +
      alpha_lb_ind[i] +
      d3 * zeros_vec[i] +
      beta_sla * scc_log_SLA[i] +
      beta_ln * scc_log_LeafN[i]


   ## Cone Cost
    logConeCost[i] ~ dnorm(
      alpha_cc +
      alpha_cc_sp[Species[i]] +
      alpha_cc_si[Sites[i]] +
      d4 * zeros_vec[i] +
      beta_cm * scc_log_CM[i],
      tau_cc)
    
    ## Seed Cost
    logSeedCost[i] ~ dnorm(
      alpha_sc +
      alpha_sc_sp[Species[i]] +
      alpha_sc_si[Sites[i]] +
      d5 * zeros_vec[i] +
      beta_smn * scc_log_SeedNM[i],
      tau_sc)

    
    
    ## Serotiny Benefit
    SerotinyBenefit[i] <- ilogit(logitSerotinyBenefit[i])
    logitSerotinyBenefit[i] ~ dnorm(
      alpha_serb +
      alpha_serb_sp[Species[i]] +
      alpha_serb_site[Sites[i]] +
      d7 * zeros_vec[i] +
      beta_cmserb * scc_log_CM[i],
      tau_serb)


    ## Serotiny Cost
    logSerCost[i] ~ dnorm(
      alpha_serc +
      alpha_serc_sp[Species[i]] +
      alpha_serc_si[Sites[i]] +
      d6 * zeros_vec[i] +
      beta_cmserc * scc_log_CM[i],
      tau_sercost)
    


    ## Cone Defence Benefit
    logitConeBenefit[i] <-
      alpha_cb +
      alpha_cb_sp[Species[i]] +
      alpha_cb_site[Sites[i]] +
      alpha_cb_ind[i] +
      beta_cmcb * scc_log_CM[i] +
      beta_sncb * scc_log_SeedNM[i]


    
    ## Observation model for Cone Maintenance (see below under 'Cone Maintenence')
    NClosedConesObs[Ntimesteps[i],i] ~ dpois(NClosedCones[Ntimesteps[i],i] * PropCounted[i])


    zeros_vec2[i] ~ dnorm(d1+d2+d3+d4+d5+d6+d7, 1)  # for block updating of costs/benefits, doesn't change likelihood
    
    

    # Loop through time-steps for each individual
    # NB - we use g for the time index, so as not to cause confusion with the transpose function

    for (g in 1:Ntimesteps[i]){

      ##**********##
      ## Maturity ##
      ##**********##

      Mature[g,i] ~ dbern(p_mat[g,i])
      logit(p_mat[g,i]) <- 
        a_mat +
        a_mat_sp[Species[i]] +
        a_mat_si[Sites[i]] +
        (b_mat + b_mat_sp[Species[i]]) * (logR[g,i] - 4)


    
      ##*************##
      ## Allocations ##
      ##*************##
  
      thetas[1:5,g,i] ~ ddirch(a_thetas[1:5,g,i])
      a_thetas[1:5,g,i] <- c(1, 1, Mature[g,i], Mature[g,i], Mature[g,i]) + 10^-200

      th_w[g,i] <- thetas[1,g,i]
      th_l[g,i] <- thetas[2,g,i]
      th_c[g,i] <- thetas[3,g,i]
      th_s[g,i] <- thetas[4,g,i]
      th_m[g,i] <- thetas[5,g,i]
    


      ##*****************##
      ## Cone production ##
      ##*****************##
    
      # Observation models for Cone Number
      ConeNumObs[g,i] ~ dpois(ConeNum[g,i] * PropCounted[i])


      # Calculate number of cones produced from resources allocated to cones and cone cost
      log(ConeNum[g,i]) <- log(th_c[g,i]) + logR[g,i] - logConeCost[i]


    
      ##*****************##
      ## Seed production ##
      ##*****************##

      # Whole-plant seed number
      log(NSeedsProdTot[g,i]) <- log(th_s[g,i]) + logR[g,i] - logSeedCost[i]


      ##**************##
      ## Cone defence ##
      ##**************##
      
      # Observation model for cone defence
      DefenceObs[g,i] ~ dbin(p_def[g,i], 20)

      # expected defence probability, based on Cone Defence Benefit and site and cone ages
      logit(p_def[g,i]) <- 
        logitConeBenefit[i] +
        beta_sa * scc_log_sa[g,i] +
        beta_ca * scc_log_ca[g,i]
      
      
      # Observation model for number of seeds observed per cone
      NSeedsObs[g,i] ~ dpois(lambda_nso[g,i])
      lambda_nso[g,i] <- DefenceObs[g,i] / 20 * NSeedsProdTot[g,i] / ConeNum[g,i] + 10^-6    # tiny offset added to ensure lambda_nso > 0


      ##******************##
      ## Cone Maintenance ##
      ##******************##
      
      # Calculate the number of cones to care for, based on resources allocated and Serotiny Cost
      log(NConesCaredFor[g,i]) <- log(th_m[g,i]) + logR[g,i] - log(delta_t[i]) - logSerCost[i]

      # Calulate number of cones remaining closed on the whole plant until the next time-step
      NClosedCones[g,i] <- (SerotinyBenefit[i] ^ delta_t[i]) * NConesCaredFor[g,i]


      # observation model for age-stratified closed cone number:
      NClosedConesObs_str[g,i] ~ dbinom( SerotinyBenefit[i] ^ ((Ntimesteps[i]-g+1)*delta_t[i]) , ConeNumObs[g,i] )
      ## NB - see above for observation model for whole-plant cone maintenance, outside of the time loop


      ##*****************##
      ## Wood production ##
      ##*****************##

      # NB - we work with basal diameter as the working variable, rather than basal area,
      # as this is more numerically stable.
      # Allometry parameters and data are thus based on the diameter:SSL relationship, 
      # then basal diameter is converted to area 


      # First, allow the basal-diameter : summed stem length (SSL) allometry to deviate for 
      # seedlings in the first time-step
      alpha_bd_hat[g,i] <-
        ifelse(g==1,
               alpha_bd[i] * gamma_alpha_bd[Species[i]],
               alpha_bd[i])

      # Calculate SSL from resources allocated to wood, basal-diameter : SSL allometry and wood density
      SSL[g+1,i] <- ( ( ( th_w[g,i] * exp(logR[g,i]) )
                          / ((alpha_bd_hat[g,i])^2 * WD[i] * (PI/4) )
                          + (SSL[g,i])^(2*beta_bd[i] + 1) ) ^ (1 / (2*beta_bd[i] + 1)))

      # Expected stem length increment, SL
      SL[g,i] <- SSL[g+1,i] - SSL[g,i] + 10^-6


      # For some plants at early ages, SL was missing data, and we only knew the SSL up that time-point.
      # As growth for these plants if fairly regular (i.e. don't expect to see 0.1cm growth followed by 10cm),
      # we thus place a very soft constraint on the missing SL measures. We say that across the time-steps with missing SLs,
      # these SLs should approximately equal SSL for this period divided by the number of time-steps,
      # with variance of the square root of this expected value
      mu_missing_SL[g,i] ~ dnorm(SL[g,i], tau_missing_SL[g,i])


      # Observation models for SL

      # Variance of stem length increments across a cohort is approximately equal to the expected SL
      # (but for the first two time-steps (when no branching occured), set this to 0.1)
      var_sl[g,i] <- (g<3) * 0.1 + (g>2) * SL[g,i]

      SLobs[g,i] ~ dnorm(SL[g,i], 1/var_sl[g,i]) T(0,)

      # ensure that SLs sum up to SSL
      # we allow very small variance for this node, to facilitate mixing
      SSLobs[g+1,i] ~ dnorm(SLobs[g,i] + SSLobs[g,i], 11)


      ##*****************##
      ## Leaf production ##
      ##*****************##

      # Calculate basal area, BA, from SSL and allometry of diameter : SSL
      logBA[g,i] <- log(PI/4) + 2*log(alpha_bd_hat[g,i]) + 2*beta_bd[i]*log(SSL[g+1,i])


      # Calculate leaf mass produced from resources allocated to leaves and leaf cost
      logLProd[g,i] <- log(th_l[g,i]) +
        logR[g,i] -
        logLeafCost[i]


      # Allow the allometry between leaf mass produced and BA to vary for seedlings in the first time-step
      lm_ba_allom_dev[g,i] <-
        ifelse(g==1,
               gamma_lm_ba_allom_sp[Species[i]] + gamma_lm_ba_allom_si[Sites[i]],
               0)

      log_alpha_lm_ba_mat[g,i] ~ dnorm(mu_laltm[g,i], tau_lm)
      mu_laltm[g,i] <-
        logLProd[g,i] -
        lm_ba_allom_dev[g,i] -
        logBA[g,i]


      ##*****************##
      ## Total Leaf Mass ##
      ##*****************##
      
      for (coh in g:(Ntimesteps[i])){
        LRem[g,coh,i] <- exp( - ( delta_t[i]*(coh-g)/beta_lrem[i] ) ^ alpha_lrem[i] ) * exp(logLProd[g,i])
      }

      TLM[g,i] <- sum(LRem[1:g,g,i])


      ##**********************##
      ## Resource acquisition ##
      ##**********************##

      # Resource Pool in the next time-step, based on Leaf Benefit, Total Leaf Mass and process error
      logR[g+1,i] ~ dnorm(logLeafBenefit[i] + log(delta_t[i]) + log(TLM[g,i]), tau_R)


    }  # close time-steps loop

  }  # close individuals loop
  
  
  


  ##*********************##
  ## PRIOR DISTRIBUTIONS ##
  ##*********************##
  # All priors are given weakly informative distributions (i.e. placing the bulk of the probability density in biologically plausible regions), 
  # except for the effect of seed N content on seed benefit, informed by a large seedling experiment comprising the same species (Cooksley et al. 2023),
  # and the residual error of leaf mass produced per branch area, informed by our empirical fit (see 'Reconstructing whole-plant vegetative growth' in the main text)


  ##*************************##
  ## Priors for seed benefit ##
  ##*************************##

  alpha_sb   ~ dnorm(0,0.01)
  beta_snmsb  ~ dnorm(0.6,20)
  sigma_r1   ~ dgamma(1,0.1)
  tau_r1    <- 1/(sigma_r1^2)
  
  for (s in 1:NSpecies){
    alpha_sb_sp[s] ~ dnorm(0, tau_sb_sp)
  }
  sigma_sb_sp ~ dgamma(2,2.5)
  tau_sb_sp <- 1/(sigma_sb_sp^2)

  for (si in 1:NSites){
    alpha_sb_site[si] ~ dnorm(0, tau_sb_site)
    beta_snmsb_si[si] ~ dnorm(0, tau_beta_snmsb_si)
  }
  sigma_sb_site ~ dgamma(2,2.5)
  tau_sb_site <- 1/(sigma_sb_site^2)
  sigma_beta_snmsb_si ~ dgamma(2,2.5)
  tau_beta_snmsb_si <- 1/(sigma_beta_snmsb_si^2)


  ##*********************##
  ## Priors for maturity ##
  ##*********************##

  a_mat1   ~ dnorm(0,0.0025)
  a_mat2   ~ dgamma(2,1)
  b_mat1    ~ dnorm(0,0.1)
  b_mat2    ~ dgamma(2,1)
  a_mat <- a_mat1 * a_mat2
  b_mat <- b_mat1 * b_mat2
  
  for (s in 1:NSpecies){
    a_mat_sp[s] ~ dnorm(0, tau_a_mat_sp)
    b_mat_sp[s] ~ dnorm(0, tau_b_mat_sp)
  }
  sigma_a_mat_sp ~ dgamma(1,0.1)
  tau_a_mat_sp <- 1/(sigma_a_mat_sp^2)
  sigma_b_mat_sp ~ dgamma(1,0.1)
  tau_b_mat_sp <- 1/(sigma_b_mat_sp^2)

  for (si in 1:NSites){
    a_mat_si[si] ~ dnorm(0, tau_a_mat_si)
  }
  sigma_a_mat_si ~ dgamma(1,0.1)
  tau_a_mat_si <- 1/(sigma_a_mat_si^2)


  ##**********************##
  ## Priors for cone cost ##
  ##**********************##

  alpha_cc   ~ dnorm(0,0.001) 
  beta_cm    ~ dnorm(0,0.01)
  
  for (s in 1:NSpecies){
    alpha_cc_sp[s] ~ dnorm(0, tau_alpha_cc_sp)
  }
  sigma_alpha_cc_sp ~ dgamma(1,0.1)
  tau_alpha_cc_sp <- 1/(sigma_alpha_cc_sp^2)

  for (si in 1:NSites){
    alpha_cc_si[si] ~ dnorm(0, tau_alpha_cc_si)
  }
  sigma_alpha_cc_si ~ dgamma(1,0.1)
  tau_alpha_cc_si <- 1/(sigma_alpha_cc_si^2)


  sigma_cc ~ dgamma(5,12)
  tau_cc <- 1/(sigma_cc^2)

  
  
  ##**********************##
  ## Priors for seed cost ##
  ##**********************##
  
  alpha_sc    ~ dnorm(0,0.001)
  beta_smn    ~ dnorm(0,0.01)
  
  for (s in 1:NSpecies){
    alpha_sc_sp[s] ~ dnorm(0, tau_alpha_sc_sp)
  }
  sigma_alpha_sc_sp ~ dgamma(1,0.1)
  tau_alpha_sc_sp <- 1/(sigma_alpha_sc_sp^2)

  for (si in 1:NSites){
    alpha_sc_si[si] ~ dnorm(0, tau_alpha_sc_si)
  }
  sigma_alpha_sc_si ~ dgamma(1,0.1)
  tau_alpha_sc_si <- 1/(sigma_alpha_sc_si^2)

  sigma_sc ~ dgamma(1,0.1)
  tau_sc <- 1/(sigma_sc^2)


  ##*****************************##
  ## Priors for serotiny benefit ##
  ##*****************************##

  alpha_serb ~ dnorm(0,0.01)
  beta_cmserb  ~ dnorm(0,0.01)
  sigma_serb1 ~ dnorm(0,1) T(0,)      # parameter-expanded to facilitate mixing
  sigma_serb2 ~ dgamma(1.5,0.1)
  sigma_serb <- sigma_serb1 * sigma_serb2
  tau_serb <- 1/(sigma_serb^2)

  for (s in 1:NSpecies){
    alpha_serb_sp[s] ~ dnorm(0, tau_alpha_serb_sp)
  }
  sigma_alpha_serb_sp ~ dgamma(1,0.5)
  tau_alpha_serb_sp <- 1/(sigma_alpha_serb_sp^2)

  for (si in 1:NSites){
    alpha_serb_site[si] ~ dnorm(0, tau_alpha_serb_site)
  }
  sigma_alpha_serb_site ~ dgamma(1,0.5)
  tau_alpha_serb_site <- 1/(sigma_alpha_serb_site^2)

  
  
  ##**************************##
  ## Priors for serotiny cost ##
  ##**************************##

  alpha_serc ~ dnorm(0,0.01)
  beta_cmserc ~ dnorm(0,0.01) 
  sigma_sercost ~ dgamma(4,10)
  tau_sercost <- 1/(sigma_sercost^2)
  
  for (s in 1:NSpecies){
    alpha_serc_sp[s] ~ dnorm(0, tau_a_serc_sp)
  }
  sigma_a_serc_sp ~ dgamma(1,0.5)
  tau_a_serc_sp <- 1/(sigma_a_serc_sp^2)

  for (si in 1:NSites){
    alpha_serc_si[si] ~ dnorm(0, tau_a_serc_si)
  }
  sigma_a_serc_si ~ dgamma(1,0.5)
  tau_a_serc_si <- 1/(sigma_a_serc_si^2)


  ##*****************************************************************************##
  ## Priors for cone defence benefit and effect of site and cone ages on defence ##
  ##*****************************************************************************##

  alpha_cb ~ dnorm(0,0.1)
  beta_cmcb ~ dnorm(0,0.2)
  beta_sncb ~ dnorm(0,0.2)
  beta_sa  ~ dnorm(0,0.2)
  beta_ca  ~ dnorm(0,0.2)
  
  for (s in 1:NSpecies){
    alpha_cb_sp[s] ~ dnorm(0, tau_cb_sp)
  }
  sigma_cb_sp ~ dgamma(1,0.1)
  tau_cb_sp <- 1/(sigma_cb_sp^2)

  for (si in 1:NSites){
    alpha_cb_site[si] ~ dnorm(0, tau_cb_site)
  }
  sigma_cb_site ~ dgamma(1,0.1)
  tau_cb_site <- 1/(sigma_cb_site^2)

  for (ind in 1:NInds){
    alpha_cb_ind[ind] ~ dnorm(0, tau_cb_ind)
  }
  sigma_cb_ind ~ dgamma(1,0.1)
  tau_cb_ind <- 1/(sigma_cb_ind^2)


  ##**********************************************##
  ## Priors for allometry variation for seedlings ##
  ##**********************************************##

  mu_gamma_alpha_bd ~ dnorm(1,1)
  for (s in 1:NSpecies){
    gamma_alpha_bd[s] ~ dnorm(mu_gamma_alpha_bd, tau_mu_gamma_alpha_bd) T(0,)
  }
  sigma_mu_gamma_alpha_bd ~ dgamma(1,1)
  tau_mu_gamma_alpha_bd <- 1/(sigma_mu_gamma_alpha_bd^2)


  mu_gamma_lm_ba_allom ~ dnorm(0,0.1)
  for (s in 1:NSpecies){
    gamma_lm_ba_allom_sp[s] ~ dnorm(mu_gamma_lm_ba_allom, tau_gamma_lm_ba_allom_sp)
  }
  sigma_gamma_lm_ba_allom_sp ~ dgamma(1,1)
  tau_gamma_lm_ba_allom_sp <- 1/(sigma_gamma_lm_ba_allom_sp^2)

  for (si in 1:NSites){
    gamma_lm_ba_allom_si[si] ~ dnorm(0, tau_gamma_lm_ba_allom_si)
  }
  sigma_gamma_lm_ba_allom_si ~ dgamma(1,1)
  tau_gamma_lm_ba_allom_si <- 1/(sigma_gamma_lm_ba_allom_si^2)


  ##************************************************************##
  ## Priors for leaf cost and benefit, and resource acquisition ##
  ##************************************************************##

  alpha_lc  ~ dnorm(0,0.01)
  beta_csla1  ~ dnorm(0,0.04)
  beta_csla2  ~ dnorm(0,0.04)
  beta_csla  <- beta_csla1 + beta_csla2    # parameter-expanded to facilitate mixing
  beta_cln1  ~ dnorm(0,0.04)
  beta_cln2  ~ dnorm(0,0.04)
  beta_cln  <- beta_cln1 + beta_cln2       # parameter-expanded to facilitate mixing
  
  alpha_lb   ~ dnorm(0,0.01)
  beta_sla  ~ dnorm(0,0.01)
  beta_ln ~ dnorm(0,0.01)

  for (s in 1:NSpecies){
    alpha_lc_sp[s] ~ dnorm(0,tau_alpha_lc_sp)
  }
  sigma_alpha_lc_sp ~ dgamma(1.2,2)
  tau_alpha_lc_sp <- 1/(sigma_alpha_lc_sp^2)

  for (si in 1:NSites){
    alpha_lc_si[si] ~ dnorm(0,tau_alpha_lc_si)
  }
  sigma_alpha_lc_si ~ dgamma(1.2,2)
  tau_alpha_lc_si <- 1/(sigma_alpha_lc_si^2)

  sigma_lc ~ dgamma(4,12)
  tau_lc <- 1/(sigma_lc^2)
  
  
  sigma_lm ~ dgamma(120,780)    # dgamma(124,782)
  tau_lm <- 1/(sigma_lm^2)

  sigma_R ~ dgamma(1.2,2)
  tau_R <- 1/(sigma_R^2)

  for (s in 1:NSpecies){
    alpha_lb_sp[s] ~ dnorm(0, tau_lb_sp)
  }
  sigma_lb_sp ~ dgamma(1,0.1)
  tau_lb_sp <- 1/(sigma_lb_sp^2)

  for (si in 1:NSites){
    alpha_lb_si[si] ~ dnorm(0, tau_lb_site)
  }
  sigma_lb_si ~ dgamma(1,0.1)
  tau_lb_site <- 1/(sigma_lb_si^2)

  for (ind in 1:NInds){
    alpha_lb_ind[ind] ~ dnorm(0, tau_lb_ind)
  }
  sigma_lb_ind ~ dgamma(1,0.1)
  tau_lb_ind <- 1/(sigma_lb_ind^2)


  ##**********************************************************##
  ## Priors for dummy parameters for the block updating trick ##
  ##**********************************************************##

  d1 ~ dnorm(0,1)
  d2 ~ dnorm(0,1)
  d3 ~ dnorm(0,1)
  d4 ~ dnorm(0,1)
  d5 ~ dnorm(0,1)
  d6 ~ dnorm(0,1)
  d7 ~ dnorm(0,1)


  ##**********************************##
  ## Priors resource growth from seed ##
  ##**********************************##

  log_b_pa ~ dnorm(0,1.5)

  for (s in 1:NSpecies){
    log_b_pa_sp_hat[s] ~ dnorm(0, 1)                           # non-centred formulation of random effects
    log_b_pa_sp[s] <- log_b_pa_sp_hat[s] * sigma_log_b_pa_sp
  }
  sigma_log_b_pa_sp ~ dgamma(1,1)

  for (si in 1:NSites){
    log_b_pa_si[si] ~ dnorm(0, tau_log_b_pa_site)
  }
  sigma_log_b_pa_si ~ dgamma(1,1)
  tau_log_b_pa_site <- 1/(sigma_log_b_pa_si^2)


}
", fill=TRUE, file="TRL_model.txt")


### function to set initial values of nodes
inits<-function(){list(
  alpha_lc=runif(1,-1,0),
  beta_csla1=runif(1,0,1),
  beta_csla2=runif(1,1,1.1),
  beta_cln1=runif(1,0,1),
  beta_cln2=runif(1,1,1.1),
  sigma_lc=runif(1,0.1,0.5),
  sigma_alpha_lc_sp=runif(1,0.1,1),
  sigma_alpha_lc_si=runif(1,0.1,1),
  
  alpha_cc=runif(1,0,2),
  beta_cm=runif(1,0.01,1),
  sigma_alpha_cc_sp=runif(1,0.1,1),
  sigma_alpha_cc_si=runif(1,0.1,1),
  sigma_cc=runif(1,0.2,1),
  
  alpha_sc=runif(1,0,2),
  beta_smn=runif(1,0.1,1),
  sigma_alpha_sc_sp=runif(1,0.1,1),
  sigma_alpha_sc_si=runif(1,0.1,1),
  sigma_sc=runif(1,0.1,1),
  
  alpha_serc=runif(1,0,2),
  beta_cmserc=runif(1,0.1,1),
  sigma_a_serc_sp=runif(1,0.1,1),
  sigma_a_serc_si=runif(1,0.1,1),
  sigma_sercost=runif(1,0.1,1),
  
  alpha_lb=runif(1,0,1),
  beta_sla=runif(1,-1,1),
  beta_ln=runif(1,-1,1),
  sigma_lb_sp=runif(1,0.1,1),
  sigma_lb_si=runif(1,0.1,1),
  sigma_lb_ind=runif(1,0.1,1),
  sigma_R=runif(1,0.2,0.4),
  
  alpha_sb=runif(1,0.1,0.5),
  beta_snmsb=runif(1,0.1,0.5),
  sigma_r1=runif(1,0.1,1),
  sigma_sb_sp=runif(1,0.1,1),
  sigma_sb_site=runif(1,0.1,1),
  sigma_beta_snmsb_si=runif(1,0.1,1),
  log_b_pa=runif(1,-0.5,0.5),
  sigma_log_b_pa_sp=runif(1,0.01,0.1),
  sigma_log_b_pa_si=runif(1,0.01,0.1),
  
  alpha_serb=runif(1,0.1,2),
  beta_cmserb=runif(1,0.1,0.6),
  sigma_alpha_serb_sp=runif(1,0.1,1),
  sigma_alpha_serb_site=runif(1,0.1,1),
  sigma_serb1=runif(1,0.1,0.5),
  sigma_serb2=runif(1,0.1,0.5),

  alpha_cb=runif(1,-2,2),
  beta_cmcb=runif(1,0,1),
  beta_sncb=runif(1,0,1),
  beta_sa=runif(1,0.1,1),
  beta_ca=runif(1,0.1,1),
  sigma_cb_sp=runif(1,0.1,1),
  sigma_cb_site=runif(1,0.1,1),
  sigma_cb_ind=runif(1,0.1,1),
  
  a_mat1=runif(1,-0.1,0.1),
  a_mat2=runif(1,0.1,0.2),
  b_mat1=runif(1,-0.1,0.1),
  b_mat2=runif(1,0.1,0.2),
  sigma_a_mat_sp=runif(1,0.01,0.1),
  sigma_a_mat_si=runif(1,0.01,0.1),
  sigma_b_mat_sp=runif(1,0.01,0.1),
  
  mu_gamma_alpha_b=runif(1,0.1,1),
  sigma_mu_gamma_alpha_bd=runif(1,0.1,1),
  mu_gamma_lm_ba_allom=runif(1,0.1,1),
  sigma_gamma_lm_ba_allom_sp=runif(1,0.1,1),
  sigma_gamma_lm_ba_allom_si=runif(1,0.1,1),
  sigma_lm=runif(1,0.1,0.2)
)}


### parameters to monitor
parameters <- c(
  "alpha_lc",
  "beta_csla",
  "beta_cln",
  "alpha_lc_sp",
  "alpha_lc_si",
  "sigma_alpha_lc_sp",
  "sigma_alpha_lc_si",
  "sigma_lc",
  
  "alpha_cc",
  "beta_cm",
  "alpha_cc_sp",
  "alpha_cc_si",
  "sigma_alpha_cc_sp",
  "sigma_alpha_cc_si",
  "sigma_cc",
  
  "alpha_sc",
  "beta_smn",
  "alpha_sc_sp",
  "alpha_sc_si",
  "sigma_alpha_sc_sp",
  "sigma_alpha_sc_si",
  "sigma_sc",
  
  "alpha_serc",
  "beta_cmserc",
  "alpha_serc_sp",
  "alpha_serc_si",
  "sigma_a_serc_sp",
  "sigma_a_serc_si",
  "sigma_sercost",
  
  "alpha_lb",
  "beta_sla",
  "beta_ln",
  "alpha_lb_sp",
  "alpha_lb_si",
  "alpha_lb_ind",
  "sigma_lb_sp",
  "sigma_lb_si",
  "sigma_lb_ind",
  "sigma_R",
  
  "alpha_sb",
  "beta_snmsb",
  "alpha_sb_sp",
  "alpha_sb_site",
  "sigma_sb_sp",
  "sigma_sb_site",
  "beta_snmsb_si",
  "sigma_beta_snmsb_si",
  "sigma_r1",
  
  "log_b_pa",
  "log_b_pa_sp",
  "log_b_pa_si",
  "sigma_log_b_pa_sp",
  "sigma_log_b_pa_si",
  
  "alpha_serb",
  "beta_cmserb",
  "alpha_serb_sp",
  "alpha_serb_site",
  "sigma_alpha_serb_sp",
  "sigma_alpha_serb_site",
  "sigma_serb",
  
  "alpha_cb",
  "beta_cmcb",
  "beta_sncb",
  "beta_sa",
  "beta_ca",
  "alpha_cb_sp",
  "alpha_cb_site",
  "alpha_cb_ind",
  "sigma_cb_sp",
  "sigma_cb_site",
  "sigma_cb_ind",
  
  "a_mat",
  "b_mat",
  "a_mat_sp",
  "b_mat_sp",
  "a_mat_si",
  "sigma_a_mat_sp",
  "sigma_b_mat_sp",
  "sigma_a_mat_si",
  
  "mu_gamma_alpha_ba",
  "gamma_alpha_ba",
  "sigma_mu_gamma_alpha_ba",
  "log_mu_gamma_alpha_ba",
  
  "mu_gamma_lm_ba_allom",
  "gamma_lm_ba_allom_sp",
  "sigma_gamma_lm_ba_allom_sp",
  "gamma_lm_ba_allom_si",
  "sigma_gamma_lm_ba_allom_si",
  
  "sigma_lm",
  
  "th_w",
  "th_l",
  "th_c",
  "th_s",
  "th_m",
  
  "logLeafCost",
  "logCCost",
  "logSeedCost",
  "logSerCost",
  "logLeafBenefit",
  "logitConeBenefit",
  "logSeedBenefit",
  "logitSerotinyBenefit",
  
  "Mature",
  "logR",
  "SL",
  "SSL",
  "SLobs",
  "SSLobs",

  "logLProd",
  "LRem",
  "TLM",
  
  "ConeNum",
  "ConeNumObs",

  "NSeedsProdTot",
  "DefenceObs",

  "NSeedsObs",

  "NConesCaredFor",
  "NClosedCones",
  "NClosedCones_str",
)


## fit the TRL model

fit1 <- saveJAGS(data = TRL_data, 
                 inits = inits, 
                 params = parameters,
                 modelFile = "TRL_model.txt",
                 chains = 10,
                 burnin = 20000, 
                 sample2save = 10, 
                 nSaves = 200, 
                 thin = 20,
                 fileStub = "give_prefix_for_files_here")  # change as necessary

# NB - we run more chains than necessary, as setting reasonable initialial values for a non-linear model this large is challenging, 
# and some chains may fail to calculate log likelihoods at starting values.
# This is, in our opinion, much preferable to initialising chains at very narrow values, e.g. close to the posterior distribution.
# Rather, we initialise at relatively overdispersed values compared to the posterior, ensuring proper exploration of possible parameter values.
# Chains that fail to initialise properly can simply be removed form the working directory before reading samples back into R.


### read back in the saved MCMC samples

recsav <- recoverSaves("give_prefix_for_files_here")
bla <- combineSaves(recsav, thin=10, cores=6, burnFiles=0)   # change cores as necessary - allows faster readingin of samples
